# Intragenic DNA inversions expand bacterial coding capacity

**DOI:** 10.1101/2023.03.11.532203

**Authors:** Rachael B. Chanin, Patrick T. West, Ryan M. Park, Jakob Wirbel, Gabriella Z. M. Green, Arjun M. Miklos, Matthew O. Gill, Angela S. Hickey, Erin F. Brooks, Ami S. Bhatt

## Abstract

Bacterial populations that originate from a single bacterium are not strictly clonal. Often, they contain subgroups with distinct phenotypes. Bacteria can generate heterogeneity through phase variation: a preprogrammed, reversible mechanism that alters gene expression levels across a population. One well studied type of phase variation involves enzyme-mediated inversion of specific intergenic regions of genomic DNA. Frequently, these DNA inversions flip the orientation of promoters, turning ON or OFF adjacent coding regions within otherwise isogenic populations. Through this mechanism, inversion can affect fitness, survival, or group dynamics. Here, we develop and apply bioinformatic approaches to discover thousands of previously undescribed phase-variable regions in prokaryotes using long-read datasets. We identify ‘intragenic invertons’, a surprising new class of invertible elements found entirely within genes, in bacteria and archaea. To date, inversions within single genes have not been described. Intragenic invertons allow a gene to encode two or more versions of a protein by flipping a DNA sequence within the coding region, thereby increasing coding capacity without increasing genome size. We experimentally characterize specific intragenic invertons in the gut commensal *Bacteroides thetaiotaomicron*, presenting a ‘roadmap’ for investigating this new gene-diversifying phenomenon.

**One-Sentence Summary:** Intragenic DNA inversions, identified using long-read sequencing datasets, are found in many phyla across the prokaryotic tree of life.

## Introduction

Adaptation is a cornerstone of survival for any species. In the complex gut microenvironment, bacteria experience many stressors including nutritional and niche competition, oxidative and nitrosative stress, and antibiotics. To overcome these challenges, bacteria may activate specific response programs which alter transcriptional or translational profiles promoting survival during these conditions. Additionally, bacterial daughter cells may acquire mutations, such as single nucleotide variations or small insertions or deletions, within genes. These gene alterations can then promote survival in the right circumstances. For example, mutations in drug targets, efflux pumps, or their regulators can provide increased resistance to antibiotics ^1,2^. While many of these gene-varying mutations in bacteria are semi-reproducible, meaning that nucleotide alterations will occur in the same region of a genome under a similar environmental stressor, most are not reversible and may be costly when the stimulus is removed^3^.

Beyond mutations and small insertions and deletions, there are only a few known mechanisms for introducing gene variation in bacteria. These mechanisms include: alternative translational start sites or terminators, which enable the encoding of two or more different gene products from a single mRNA ^4,5^; slipped-strand mispairing, which introduces replicative or translational changes that can alter bacterial gene sequence length ^6^; and diversity generating retroelements ^7^, which can diversify a gene during reverse transcription and recombine in a novel gene variant. Outside of these rare gene-varying events, the typical prokaryotic ‘one gene, one gene product’ rule generally holds and is in stark contrast to nearly all Eukaryotes, in which a large proportion of transcribed genes can undergo alternative splicing to generate multiple protein isoforms from one gene.

One fairly prevalent mechanism of reversible adaptation in bacteria is phase variation. This is a preprogrammed and reversible mechanism that generates phenotypic diversity in a clonal population ^8^. Phase variation can promote cooperativity by sharing resources between subgroups in a metabolically efficient way or through bet hedging by diversifying a population to protect from complete elimination in future selective events. One type of preprogrammed variation occurs through DNA inversion. Site-specific recombinases recognize a pair of inverted repeats in genomic DNA and invert the intervening DNA sequence ^9^. In the first described example, DNA inversion of a promoter sequence resulted in the switching of expression from one flagellar antigen (H1) to another (H2) in *Salmonella enterica* serovar Typhimurium. The change in antigen expression determined whether the bacterium was bound by antiserum, and thus was termed a ‘phase-determining event’^10–12^. This DNA inversion, and the many others that have since been discovered, play critical adaptive roles in both commensal and pathogenic bacteria.

For decades, these invertible loci were identified individually. Then, computational approaches enabled higher-throughput discovery of these ‘invertons’ across the genomes of a small subset of specific bacterial species ^13,14^. In 2019, Jiang *et al.* developed an elegant method that facilitated broad scale identification of 4,686 intergenic invertons (i.e., invertons between genes) through a search of 54,875 bacterial reference genomes ^15^, utilizing short-read mapping as evidence; however, short-reads cannot span entire invertons, which can range in lengths of up to multiple kbps ^15^. The long length of many of these invertons causes short-read based inverton detection methods to be lower in sensitivity, as methods to detect invertons rely on reads that span one boundary of a given inverton. This means that only a small proportion of reads provide usable evidence for inverton detection. Similarly to Jiang *et al*., in early 2023, Milman *et al.* used a computational model to predict over 11,000 potential invertons that partially overlap with genes (partial intergenic) in >35,000 bacterial species. Partial intergenic invertons are sometimes referred to as shufflon systems; they function by flipping out homologous domains of enzymes, which can change their specificity ^16,17^. Once Milman *et al.* predicted the candidate invertons, they then manually inspected publicly available long-read datasets, which led to the validation of 22 of the >11,000 predicted invertons ^18^. Taken together, these two studies demonstrate that computational approaches can be a powerful method to identify invertible elements within many phyla. Furthermore, the ubiquity of both intergenic and shufflon-type invertons in bacterial genomes highlights their likely importance in affecting bacterial gene regulation and phenotypes.

While previous work has demonstrated the presence of intergenic and shufflon-type invertons, there are no reports to our knowledge of invertons that occur entirely within genes. Such invertons would represent a novel mechanism of preprogrammed gene variation in bacteria. Furthermore, with the advent of long-read sequencing technologies, and improvements in their accuracy, developing a long-read inverton finding workflow would be expected to improve sensitivity in inverton discovery and detection. Building off these concepts, here we find that the same mechanisms that underlie intergenic and partially intergenic invertons can occur entirely within a gene. These intragenic invertons expand bacterial coding capacity by either recoding protein sequences within the inverted region or introducing premature stop codons. In both cases, intragenic invertons result in a single gene being able to produce two or more different protein products. We develop PhaVa, a long-read based tool to identify intragenic, intergenic, and partial intergenic invertons. By applying PhaVa to long read sequencing data for ∼30,000 bacterial isolates from ∼4,000 unique species, we find that intragenic invertons occur in many phyla across the prokaryotic tree of life. In particular, we focus on *Bacteroides thetaiotaomicron*, a model enteric commensal, and validate 10 intragenic invertons experimentally with particular focus on the inverton contained within the thiamine biosynthesis protein *thiC*. Finally, we make both the PhaVa software package and all of the identified invertons (intragenic, intergenic, and partial intergenic) publicly available.

## Results

Most knowledge regarding bacterial genes and their regulation is based on bacteria that are studied in laboratory conditions. Because of this, invertons that provide a fitness advantage *in vivo* but may not be advantageous to fitness *in vitro* have likely been overlooked ^19–21^. We therefore hypothesized that there are currently unknown gut-relevant invertons. To test our hypothesis, we endeavored to identify invertons in metagenomic sequencing data from longitudinally collected human stool samples from 149 adult and 21 pediatric patients undergoing hematopoietic cell transplantation ^22,23^ (Fig. 1A). These samples were selected given the varying and complex environments enteric bacteria would encounter over time, with many different stressors present such as chemotherapy, antibiotic treatment, variation in food intake, and inflammation. We hypothesized that these factors might induce inverton flipping.

**Fig. 1.**
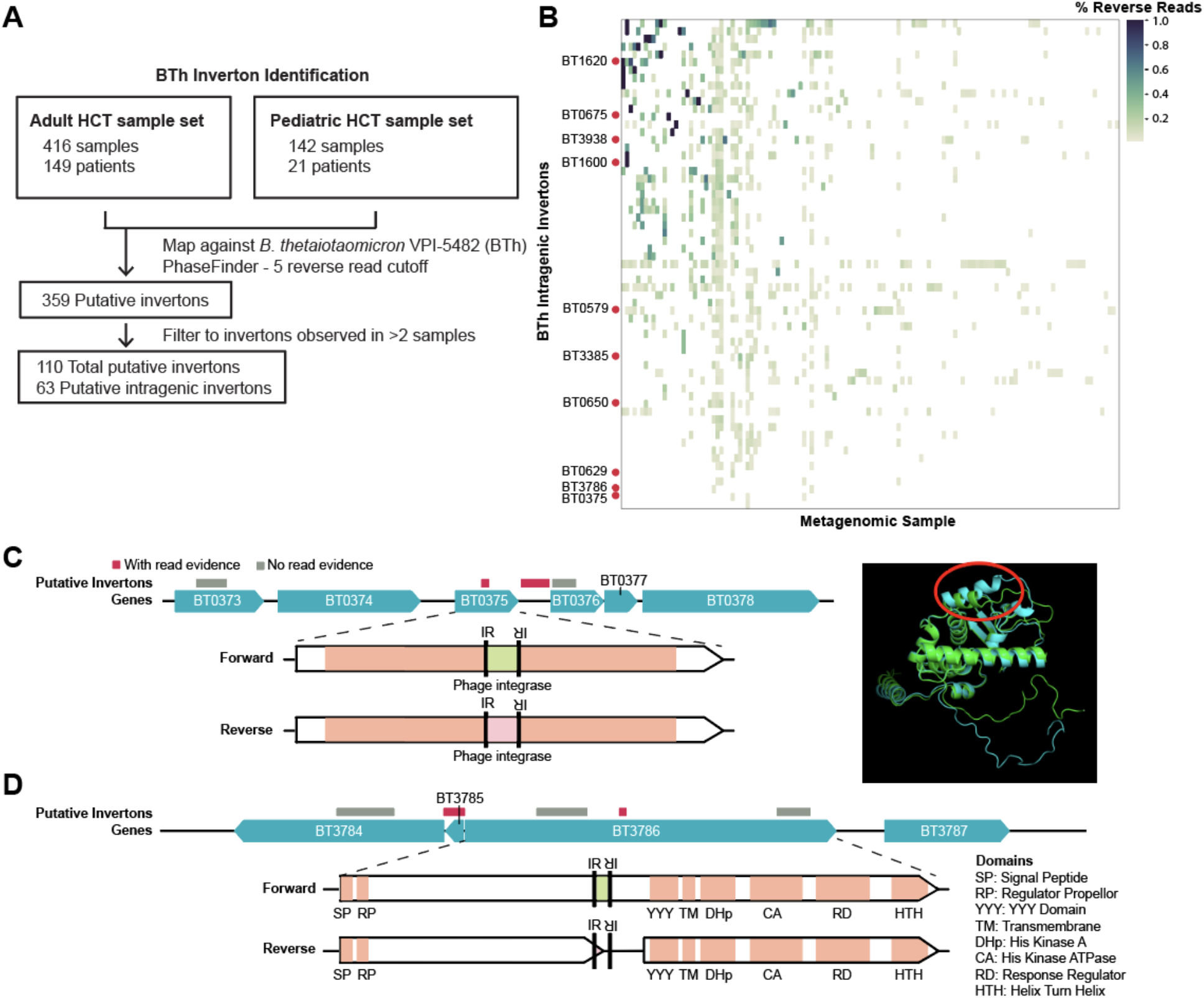
Short-read metagenomic datasets reveal intragenic invertons in *Bacteroides thetaiotaomicron* (BTh) (**A**) An overview of the analysis pipeline for identifying putative invertons in short-read datasets. (**B**) A heatmap of the inversion proportion of intragenic invertons in BTh. Samples with no intragenic invertons were removed. Rows labeled with a gene name represent intragenic invertons with PCR and Sanger sequencing evidence of inversion. (**C**-**D**) Genome diagrams for confirmed intragenic invertons in BTh. Gray bars indicate putative invertons without sequencing support. Red bars indicate invertons with sequencing evidence. (**C**) Left - Genome diagram of the region surrounding the BT0375 recoding intragenic inverton, and a domain diagram of the BT0375 gene with the location of the inverton IRs indicated. Right - AlphaFold overlay of the BT0375 forward (blue) pLDDT 89.91 and reverse (green) pLDDT 85.24. The region that is recoded is circled in red. (**D**) Genome diagram of the region surrounding the BT3786 premature stop codon intragenic inverton, and a domain diagram of the gene. The consequence of the inversion and resulting two predicted ORFs are indicated in the domain diagram.

In our efforts to comprehensively annotate invertons from this metagenomic data set, in which there are many different organisms represented in each sample, we first decided to examine invertons in organisms within the taxon Bacteroidetes. Bacteroidetes species are prevalent and typically highly abundant in the human gut, and many organisms within this taxon have known intergenic invertons ^15^. To orthogonally confirm sequencing-based observations in subsequent microbiological and genetic experiments, we focused our analysis on *B. thetaiotaomicron*, a genetically tractable species suitable for downstream experimental manipulation. To identify invertons in *B. thetaiotaomicron*, we used PhaseFinder ^15^, a short-read, reference-based inverton detection pipeline with *B. thetaiotaomicron* VPI-5482 (BTh) as the reference genome, and with relaxed filters to increase sensitivity (see methods, Fig. 1A). As an internal control to assess whether PhaseFinder could sensitively detect BTh invertons in our metagenomic samples, we examined BTh’s capsular polysaccharide (CPS) genes, a known set of invertible loci. BTh has 8 loci that encode different CPS, 5 of which are controlled by invertible promoters ^24–26^. CPS are important mediators of phage susceptibility ^27^ and can modulate the host immune system ^28–30^. Using PhaseFinder on the patient sample datasets, we found read evidence of all 5 CPS invertons in both the reference and inverted (flipped) orientations (fig. S1), demonstrating that PhaseFinder is able to detect known invertons in these metagenomic samples and that these samples have enough *Bacteroides* sequencing depth to identify invertons. Of note, in the reference BTh genome, the invertible promoter for each of these 5 loci is in the ‘OFF’ state by virtue of it being oriented in the opposite direction of the CPS genes. Similarly, *in vitro* transcriptional analyses support the finding that the majority of invertible CPS loci are in the OFF orientation ^31^, suggesting that in laboratory conditions, these loci are not transcriptionally active. Finding read evidence of inversion for all invertible CPS loci suggests that the *in vivo* patient datasets are an ideal environment to detect invertible events that are rare in laboratory-grown bacteria but may be prevalent in bacteria living in more ‘natural’ ecological settings.

In addition to known intergenic invertons such as those in the CPS loci, we also found read evidence of intragenic invertons in BTh across 132 short-read metagenomic samples (Fig. 1B). We use the term ‘intragenic inverton’ to describe invertible regions found entirely within single genes. To date, the only description of invertible DNA sequences entirely within a gene are in isolated cases of very short (7 bp) flips within mitochondrial DNA in certain pathogenic states ^32^. These 7 bp mitochondrial DNA flips are postulated to be the consequence of an enzyme-independent event, and thus are different from what we predict here to be an invertase-mediated, preprogrammed inversion. In the intragenic invertons that we observed, there were two predicted consequences. In some cases, the intragenic inverton resulted in a portion of the protein being “re-coded” (Fig. 1C). For example, we observed a 57 bp inversion in BT0375, the invertase that is believed to flip the adjacent CPS1 invertible promoter. This intragenic inversion changes the amino acid sequence of the ‘flipped’ region, and might alter the binding specificity of the invertase, possibly changing the invertible repeats (IRs) that it targets for flipping or its binding affinity for its cognate IRs. In other cases, the intragenic inverton resulted in the introduction of a ‘premature’ stop codon, affecting the prediction of protein coding open reading frames (ORFs). Often, inversion resulted in two predicted ORFs (called with Prodigal ^33^). For example, the inverton in the hybrid two-component system BT3786 occurs between two predicted protein folding domains, and thus might untether the “sensing” and “response” elements (Fig. 1D) of this signaling protein. However, we also observed intragenic inversions that resulted in zero, one, three, or more ORFs. Taken together, we describe the discovery of intragenic invertons and identify two types of invertons - those that are ‘recoding’ and those that cause a ‘premature stop’.

To validate these predicted intragenic invertons, we analyzed the DNA sequences in these gene regions *in vitro*. We extracted DNA from wild-type BTh grown in either rich or defined media and designed PCR primer sets that enabled us to amplify either the reference or the inverted version (fig. S2). We tested 59 of the 63 predicted intragenic invertons and confirmed that 10 of them had DNA molecules in both the reference and inverted orientation in our laboratory-grown population of BTh (Fig. 1B, Table 1, Data S1). The 49 unconfirmed intragenic invertons may be due to the absence of cues in the growth conditions required to flip the locus to the inverted orientation and/or false positives from the metagenomic read-based evidence.

**Table 1.** Confirmed intragenic invertons BTh. Intragenic invertons confirmed *in vitro* in BTh are listed. Invertons from short-read datasets were called with PhaseFinder on metagenomic samples (see Fig.1). The predicted consequence of inversion is also listed.

| Gene | Annotation | Consequence |
| --- | --- | --- |
| BT0375 | integrase | recoding |
| BT0579 | putative transcription regulator | premature stop codon |
| BT0629 | Mn<sup>2+</sup> and Fe<sup>2+</sup> transport protein | premature stop codon |
| BT0650 | thiamine biosynthesis protein ThiC | premature stop codon |
| BT0675 | N-acetylglucosamine-6-phosphate deacetylase NagA | recoding |
| BT1600 | BexA, membrane protein | premature stop codon |
| BT1620 | SusD homolog | premature stop codon |
| BT3385 | putative helicase | premature stop codon |
| BT3786 | two-component system sensor histidine kinase/response regulator, hybrid (one-component system) | premature stop codon |
| BT3938 | ATP-dependent DNA helicase RecQ | premature stop codon |

As genomic structural variation often involves highly repetitive or low complexity regions, short-reads are often not long enough to resolve these sequences ^34^, and thus short-read based approaches would be predicted to have limited sensitivity. We, therefore, developed a long-read based inverton predictor, PhaVa. PhaVa maps long-reads against both a forward (identical to reference) and reverse orientation version of potential invertons (Fig. 2A). PhaVa’s accuracy is high because it requires long-reads that span the entire length of a given inverton in order to make a ‘call’ about its orientation. To ensure accurate performance of PhaVa, we optimized read mapping parameters by simulating long-read datasets from ten bacterial genomes at various sequencing depths (Fig. 2B-C). The reads were generated from a reference genome, and thus no invertons are expected and any detected would be false positives. In general, the false positive rate was very low (mean false positive count per simulated sample of <0.1 in 9/10 species), with the exception of reads simulated from the *Bordetella pertussis* genome (Fig. 2D). Further investigation revealed the false positives detected in *B. pertussis* were due to a single putative inverton with inverted repeats longer than 750 bps, of which only a smaller portion of the total length were detected by ‘einverted’, the computational tool used to detected inverted repeats (fig. S3). In summary, our long-read based inverton predictor, PhaVa, demonstrates high accuracy in resolving complex genomic structural variations, with only rare instances of false positives observed.

**Fig. 2.**
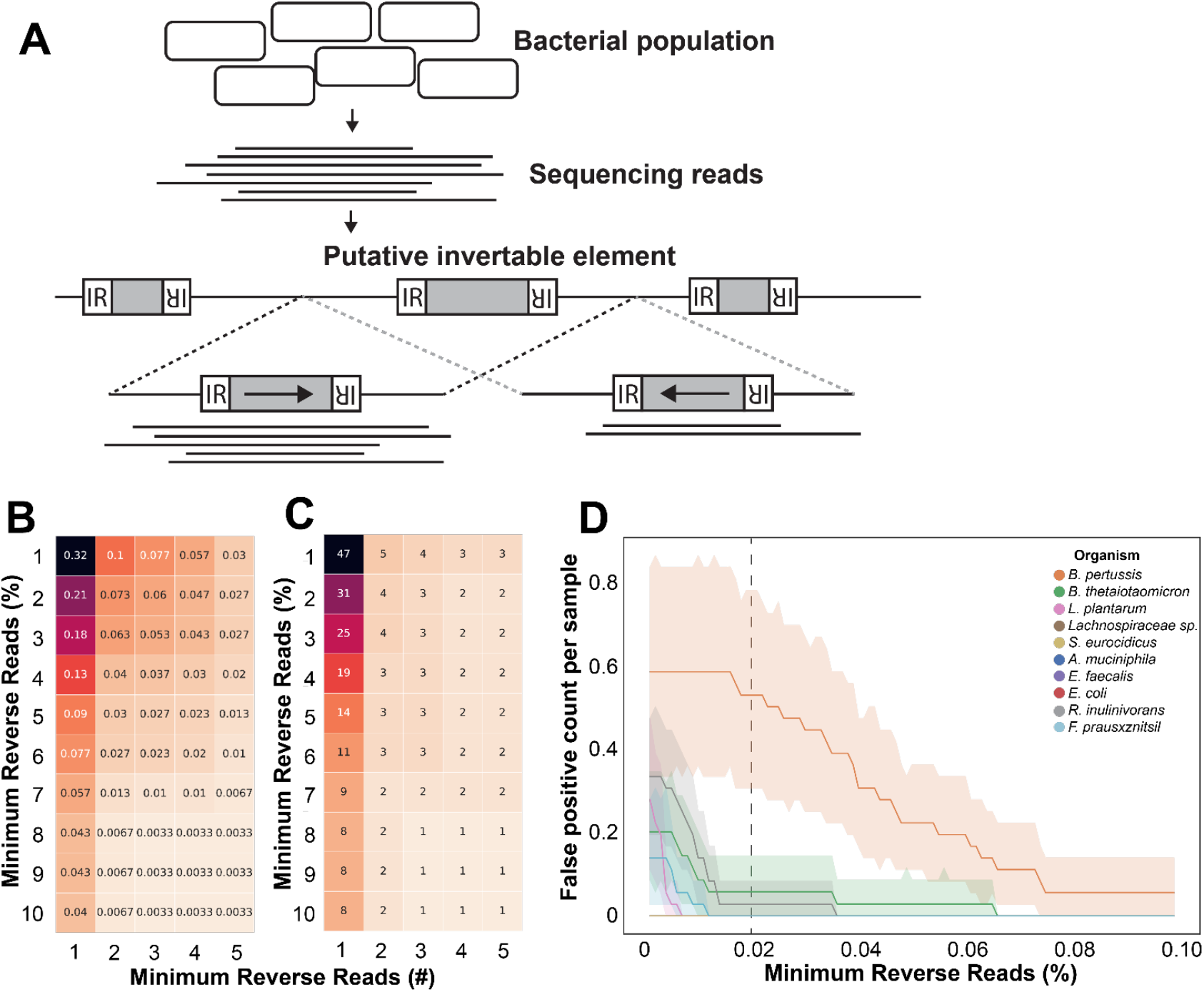
Developing and optimizing PhaVa, a long-read based, accurate inverton caller. **(A)** Schematic of PhaVa’s workflow. Putative invertons are identified, and long-reads are mapped to both a forward (highlighted by the black dashed lines) and reverse orientation (highlighted by the gray dashed lines) version of the inverton and surrounding genomic sequence, similar to PhaseFinder. Reads that do not map across the entire inverton and into the flanking sequence on either side, or have poor mapping characteristics are removed. See methods for details. **(B-C)** Optimizing cutoffs for the minimum number of reverse reads as both a raw number and percentage of all reads, to reduce false positive inverton calls with simulated reads. Cell color and number represent **(B)** the false positive rate per simulated readset and **(C)** the total number of unique false positives across all simulated datasets. **(D)** False positives in simulated data plotted per species. All measurements were made with a minimum of three reverse reads cutoff and varying the percentage of minimum reverse reads cutoff. Dashed line indicates the minimum reverse reads percent cutoff used for isolate and metagenomic datasets.

To find invertons across prokaryotic genomes, we ran PhaVa on ∼30,000 prokaryotic isolate long-read datasets deposited on SRA. We limited our analysis to readsets belonging to Bacteria or Archaea and with 50 Mbp or more of total sequencing, which resulted in our final analysis containing results from ∼4,000 unique species (fig. S4). The vast majority of these datasets represented bacteria, with only 42 archaeal long-read sequencing datasets. In total, we identified 4622 unique invertons, 3,468 of which are intergenic. Of note, compared to Jiang *et al.* ^15^, we find invertons at a higher rate per sequencing dataset (0.15 vs 0.07) and per individual isolate (1.15 v 0.09) highlighting the increased sensitivity of long-reads for detecting this type of structural variation. Like Jiang *et al.* ^15^, we found that Bacteroidetes have a relatively large number of intergenic invertons (673, Fig. 3A) and intergenic invertons per genome (2.26, Fig. 3B). Fusobacteria, Gammaproteobacteria, and Verrucomicrobia also have high numbers of intergenic invertons per genome (Fig. 3B), with Verrucomicrobia having the highest number per genome overall at 5.55 intergenic invertons per genome. In our dataset, Verrucomicrobia is composed of only *Akkermansia* strains. As increases in *Akkermansia* abundance correlate with protection against metabolic disease ^35,36^, there is interest in its use as a probiotic. However, *Akkermansia* strains exhibit broad phenotypic diversity and differential gut colonization ability ^37^, which may be attributable, in part, to the orientation of these varied intergenic invertons. In addition to the intergenic invertons, we also identified 733 partial intergenic invertons (Fig. 3A). Many of these partial intergenic invertons may form shufflon systems, and thus, as expected, these invertons are significantly longer than intergenic or intragenic invertons (Fig. 3C, p=7.1e-293 and p=7.6e-67 with a t-test, respectively). This finding of 733 partial intergenic invertons adds to the 22 long-read-validated intergenic invertons recently reported by Milman *et al*. Thus, our analysis of ∼30,000 prokaryotic isolate long-read datasets from diverse species uncovered both known and novel intergenic and partial intergenic invertons, shedding light on the remarkable structural variability within prokaryotic genomes and emphasizing the heightened sensitivity of long-read sequencing in this context.

**Fig. 3.**
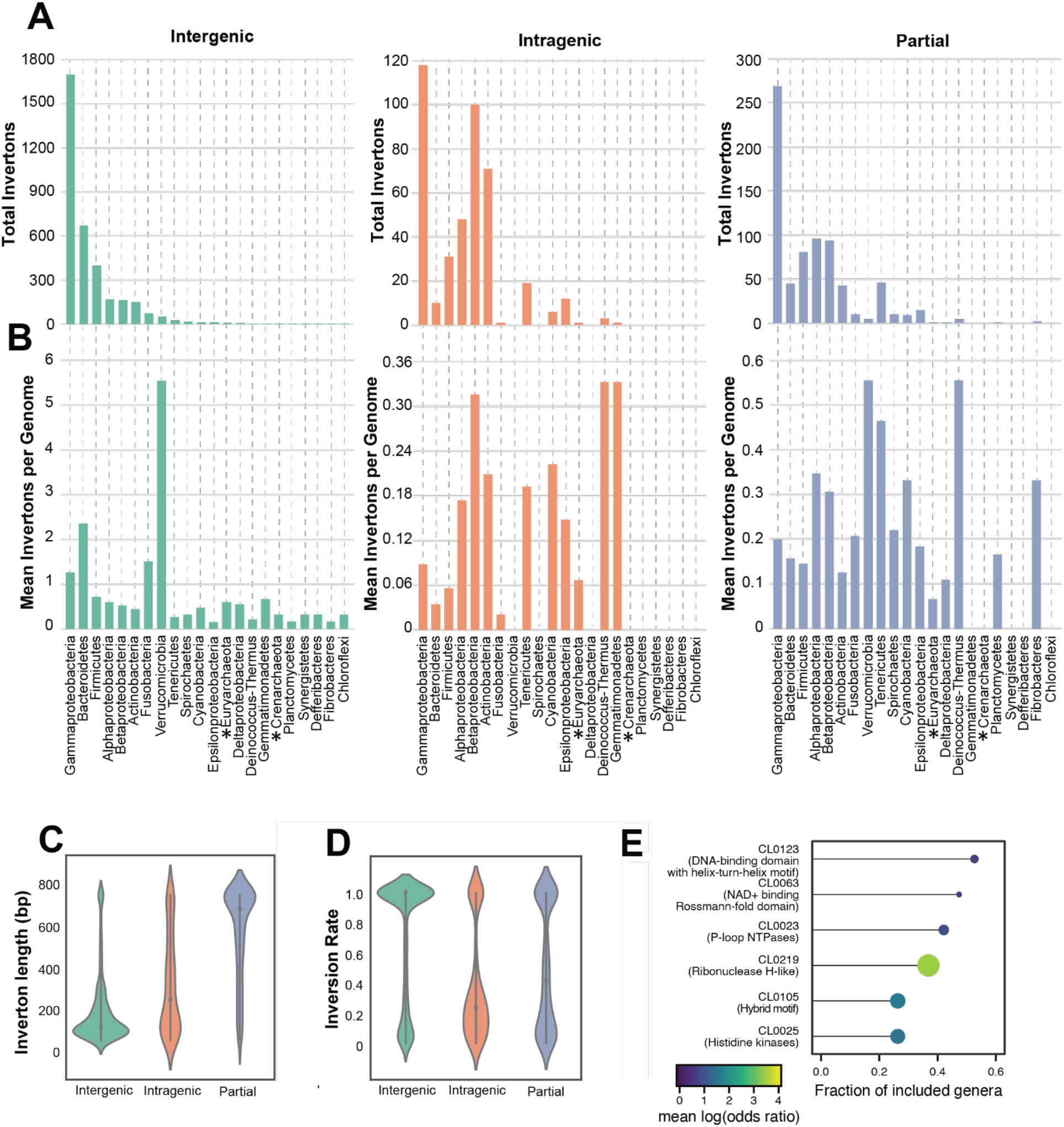
PhaVa analysis of isolate long-read sequencing data reveals intragenic inversions are prevalent across the bacterial tree of life. (**A**) The total number of invertons found within various bacterial phyla from 29,989 publicly available long-read isolate sequencing datasets. Green bars refer to intergenic invertons. Orange bars refer to intragenic invertons. Blue bars refer to partial intergenic invertons. Asterisks denote phyla within Archaea. Inset corresponds to the portion of the bar graph outlined in dotted lines. (**B**) The mean number of invertons found per genome within a phylum, of genomes that had at least one inverton. Asterisks denote phyla within Archaea. (**C**) The distribution of lengths of identified invertons, grouped by inverton type (intergenic, partial intergenic - denoted ‘partial’, and intragenic). Median value is indicated by gray dots. Partial length distribution was found to be significantly different from intergenic (p=0.0) and intragenic (p=4.5e-146) with a t-test (**D**) The distribution of inversion rates of identified invertons, defined as the percentage of reads mapped in the reverse orientation. Median value is indicated by gray dots. (**E**) Pfam clan enrichment across several genera. Dot size and fill color is proportional to the mean log-odds ratio, an effect size measure for the enrichment, and the length of the line indicates the fraction of included genera in which an enrichment score for the specific clan could be calculated.

Beyond intergenic and partial intergenic invertons, we also found evidence of intragenic invertons across multiple phyla, including the major gut microbiome-related phyla, Proteobacteria, Firmicutes and Bacteroidetes (Fig. 3A-B). We found the largest number of intragenic invertons, 118, in Gammaproteobacteria, including from organisms such as *Escherichia coli* and *Salmonella*; this is largely due to the abundance of samples for these organisms in SRA and our dataset (∼4,000 *E. coli* samples (fig. S4)), given that Gammaproteobacteria have a relatively small number of intragenic invertons detected per genome (0.09, Fig. 3B). Few long-read datasets for Archaea were available with 36 and 6 for Euryarchaeota and Crenarchaeota, respectively. Despite this, 12 putative archaeal invertons were identified; ten intergenic, one partial intergenic, and an intragenic inverton that introduces an early stop in a adenylosuccinate synthase gene in *Salarchaeum sp. JOR-1* (42 total archaeal genomes searched, Fig. 3A-B, Data S2). Chromosomal invertons have only been minimally investigated in Archaea. However, our study and a recent computational analysis of phase variable Type 1 restriction modification systems by Atack *et al.* ^17^ suggest that inverton-mediated phase variation may be an important, yet understudied, regulatory mechanism in this domain. The mean number of intragenic invertons per genome varied greatly between different phyla (Fig. 3B) with Tenericutes, Betaproteobacteria, and Actinobacteria having a relatively high number of intragenic invertons detected per genome, at 0.19, 0.32, and 0.21, respectively. The distribution of inversion proportions of individual intragenic invertons was different from that of intergenic invertons (Fig. 3D); intergenic inversions typically appeared to be in either an “ON’’ or “OFF” state in a given sample - suggesting that all of the organisms within that population shared the same biological ‘state’ of that inverton. By contrast, intragenic invertons more commonly had inversion proportions somewhere between 0 and 1 (Fig. 3D), indicating presence of both the forward and reverse orientations within a given ‘clonal’ sample. Invertons with a 100% or near 100% proportion in the ‘reverse’ orientation may also represent those that can no longer be flipped, either due to mutations in the IR or loss of the invertase that flips the inverton.

Having cataloged these intragenic invertons, we next investigated whether specific gene types or functions were enriched for the presence of intragenic invertons by doing a clade-resolved enrichment analysis. We calculated gene set enrichments (using Pfam clan definitions as gene sets) per genome, species, and genus, combining the genes from all genomes in a specific clade (Fig. 3E, S5). We found six Pfam clans enriched across several genera with the strongest and most consistent enrichments for the Pfam clans CL0123 (Helix-turn-Helix) and CL0219 (RNase-H-like) (fig. S5). This indicates that intragenic invertons occur more frequently than would be expected by chance in genes that have DNA binding or DNA/RNA modifying activity.

As noted previously, we postulate that inverton orientation likely relates to the environment of a bacterium, and that invertons are more likely to be in the non-reference orientation in organisms that are living in their ‘natural’ ecological settings. Therefore, we also ran PhaVa on 210 *de novo* assembled long-read metagenomes from the human gut ^38,39^, mapping sequencing reads back to their respective metagenomic assemblies. This enabled us to detect invertons that may be absent in isolated bacteria grown in laboratory cultures, but present *in vivo.* Doing so, we identified over 3,500 putative invertons, largely from contigs assigned to the phyla Bacteroidetes and Firmicutes (fig. S6A). In keeping with our model that invertons are more likely to be ‘active’ *in vivo* than *in vitro*, significantly more invertons were identified per species in the metagenomic samples than in the isolate sequencing samples (fig. S6B). We hypothesize this is because bacteria grown as isolates in laboratory settings do not experience the wide range of diverse environmental conditions that they do in their natural, polymicrobial habitats. Our analysis of the metagenomic data with PhaVa suggests that bioinformatic analysis of isolate genomes grown in laboratory conditions likely underestimates the number and range of invertons that exist in microbes. Therefore, the invertons called from the isolate datasets can be thought of as a ‘minimal set’, as isolate conditions may not be the ideal setting to uncover phase variable regions relative to metagenomic samples or co-cultures.

Both short-read and long-read based analyses of metagenomic datasets revealed that intragenic invertons exist. However, the biological consequences of inversion of these invertons to the non-reference orientation is not known. Thus, to evaluate the phenotypic consequences of a particularly prevalent inverton, we focused on an intragenic inverton that introduces a premature-stop codon in the BTh BT0650 gene (thiamine biosynthesis protein ThiC) (Fig. 4). Thiamine is an essential cofactor in many cellular biochemical processes and is essential for nearly all organisms. Some organisms, such as humans and certain gut microbes, are fully reliant on dietary, host, or other microbial sources for vitamins or their building blocks; others, like many gut microbes, including BTh, have the capacity to biosynthesize thiamine, albeit at a large energetic cost ^40,41^. Thus, thiamine availability has been hypothesized to strongly influence microbial community composition ^42^. We chose to characterize the intragenic inverton in *thiC* as this gene has a defined function in thiamine biosynthesis ^43,44^. Specifically, the *thiC* gene product, which encodes the enzyme 2-methyl-4-amino-5-hydroxymethylpyrimidine phosphate (HMP-P) kinase, catalyzes the conversion of aminoimidazole ribotide (AIR) to 4-amino-5-hydroxymethyl-2-methylpyrimidine (HMP) and forms a key wing in thiamine biosynthesis. In addition to having a defined role, we detected intragenic inversion in both DNA and RNA in our laboratory grown BTh strain (Fig. 4B). We predicted that the non-reference orientation of the inverton introduces a premature stop codon in the *thiC* mRNA, which would result in a truncated protein containing only the N-terminal “thiC associated domain” of the protein (Fig. 4A). The exact function of this domain of the protein is not known, but it is required for enzyme function. As noted previously, BTh can grow in the absence of exogenous thiamine as it can synthesize thiamine *de novo,* however, strains that lack ThiC lose this ability. We hypothesized that inversion of the invertible locus in *thiC* would interfere with thiamine biosynthesis, and would phenocopy the ThiC null mutant.

**Fig. 4.**
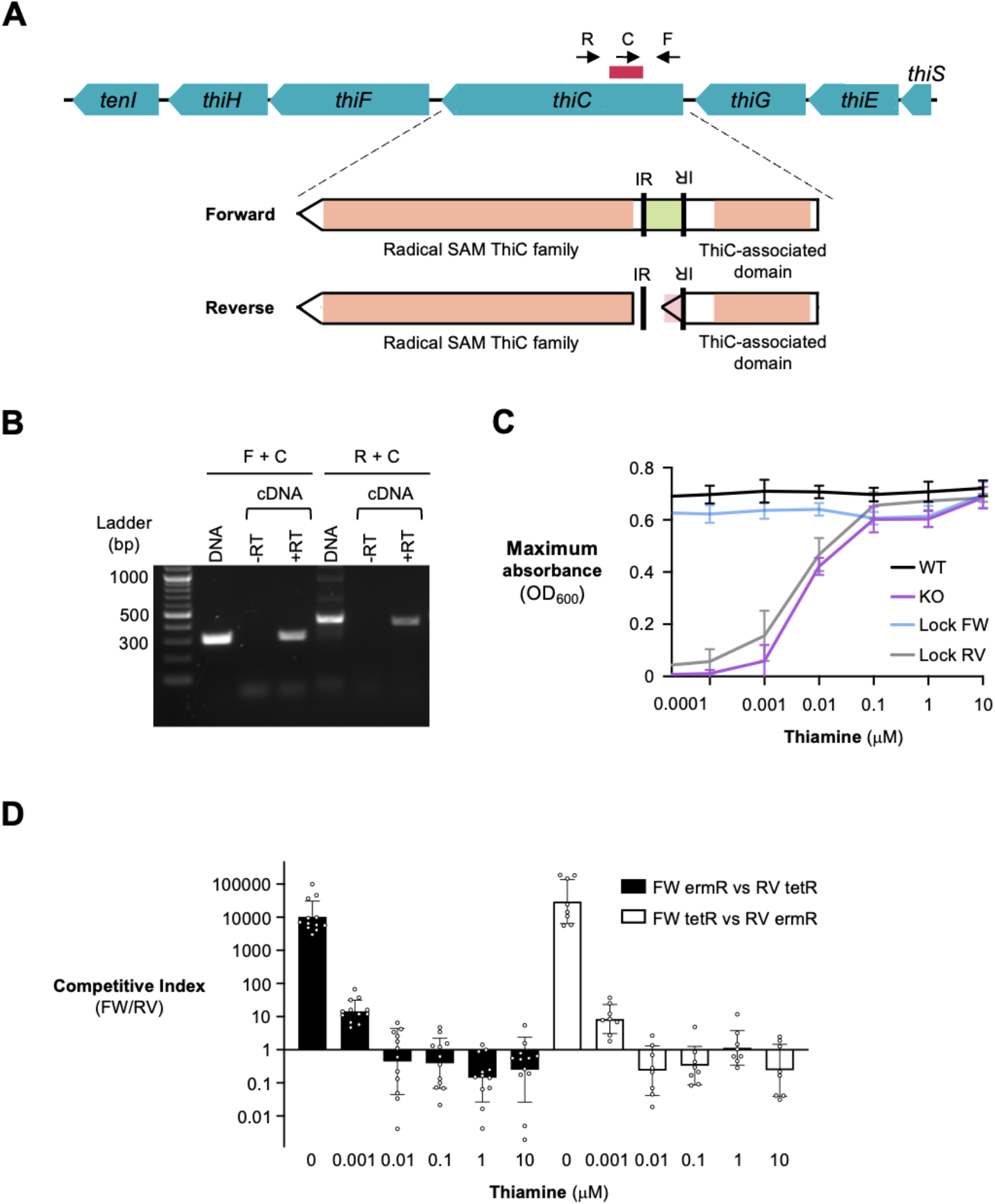
Consequences of inversion in thiamine biosynthesis protein (*thiC*) (**A**) Schematic showing the location of the *thiC* intragenic inverton (red bar). Inverton flipping results in a premature stop codon located between two protein-folding domains in ThiC. Black arrows indicate the binding location of primers used to determine the orientation of inverton. (**B**) PCR confirmation of the *thiC* intragenic inverton in both genomic DNA and reverse transcribed RNA (cDNA). PCR products of the expected size were extracted and confirmed with Sanger sequencing. (**C**) BTh strains were grown in defined media with the indicated concentrations of thiamine. The maximum optical density of each strain reached was recorded. Each point represents the average of six replicates conducted across two separate experiments. Mean and standard deviation are shown. Locked forward (blue line), locked reverse (gray line), *thiC* knockout (purple line), and wild-type (black line) are presented. (**D**) Locked strains were competed against each other in thiamine-containing media. The competitive growth experiment was performed in two different ways with the antibiotic resistance marker cassettes flipped between the two versions. Black bars indicate the locked forward strain marked with erythromycin resistance and locked reverse strain marked with tetracycline resistance. White bars indicate the locked forward strain marked with tetracycline resistance and locked reverse strain marked with erythromycin resistance. The competitive index was determined. Geometric mean and geometric standard deviation are shown for 8-12 replicates across 4-6 independent experiments.

To test the biological consequences of inversion, we generated ‘locked’ versions of the *thiC* inverton that prevent inversion from occurring within the gene. Traditionally, locking elements in a specific orientation is accomplished by mutating the nucleotides in the inverted repeat regions required for inversion or by deleting the inverted sequences entirely. Unfortunately, for intragenic invertons, deletion of these sequences or complete mutations would alter the corresponding amino acid sequences and confound interpretation. We therefore exploited the wobble position of the codon to maximize mismatches between the inverted repeats. By mutating these residues, we introduced mismatches in 6 out of 11 positions of the inverted repeat (fig. S7A). Using this method, we created a locked forward (reference orientation) and a locked reverse (flipped intragenic inverton) *thiC* strain. We also generated a *thiC* clean deletion strain.

Next, we grew wild-type BTh, locked forward, locked reverse, and the *thiC* knockout strain in various concentrations of thiamine (Fig. 4C). The locked forward strain phenocopied the wild-type strain, as it was able to grow to the same optical density regardless of whether thiamine was added to the media. By contrast, the locked reverse strain mirrored the *thiC* knock-out strain and was only able to grow to wild-type levels when 0.1 *μ*M or greater thiamine was added to the media. This finding confirms our expectation that the reverse version of the intragenic inverton interferes with ThiC function.

Having found that the locked reverse strain of the *thiC* intragenic inverton phenocopies the null mutant, we wondered whether there may be physiological circumstances that favor this mutant over the wild-type or locked forward strain. A classical approach to assess the relative fitness of two bacterial strains is to perform a competitive growth experiment. Thus, to test whether the inverted form of the *thiC* inverton provides a fitness advantage in different conditions, we competed the locked forward strain against the locked reverse strain in an equal proportion in varying concentrations of thiamine. Each strain was chromosomally marked with a different antibiotic resistance cassette. Then we determined the competitive index (CI), which is the ratio of recovered locked forward bacteria to recovered locked reverse bacteria (Fig. 4D). To account for any fitness advantages conferred by the antibiotic resistance cassettes, we repeated the competition with the cassette swapped between the two strains. While results varied slightly between these two complementary versions of the experiment, they were generally concordant. Specifically, we found that as thiamine concentration increases in the media the advantage conferred by the locked forward version of *thiC* was first diminished and then abolished at concentrations above 0.01 *μ*M. In one version, the locked reverse significantly outcompeted the locked forward strain at 1 and 10 *μ*M, whereas in the other the version the reverse significantly outcompeted the locked forward at 0.01, 0.1, and 10 *μ*M (fig. S7 B-C). Notably, at physiologically relevant thiamine concentrations, in the human intestine 0.02-2uM ^45^, the locked reverse strain was more fit than the locked forward strain. This finding complements previous work showing that auxotrophs have a fitness advantage in conditions containing a low level of exogenous metabolites when competing against prototrophic strains ^46^. The reversible nature of invertons would allow a subgroup to switch between phenotypes, whereas a simple loss of function mutation would not. Taken together, we find evidence of a physiologically relevant condition in which an intragenic inversion within the *thiC* gene may provide an energetic or other form of competitive advantage, and thus might be adaptive.

## Discussion

Bacterial genomes densely encode functional genetic programs as well as multiple layers of bioregulation. These layers of programming can be accessed by varying transcription, translation, or through genomic restructuring. One mechanism of genome restructuring and resultant ‘genomic plasticity’ is through enzyme-mediated DNA inversions. Such inversions can regulate transcription of specific genetic loci through the flipping of promoter orientation ^10,15,47,48^. Furthermore, DNA inversions can also regulate shufflon systems that recombine modular domains of bacterial protein-encoding genes to alter enzyme specificity ^18,49,50^. To date, entirely within-gene DNA inversions have not been described in prokaryotes. While not present in every genome, the identification of enzyme-mediated intragenic inversions is important as it represents another mechanism of genetic variation, and a way in which a single genetic locus can encode multiple genes.

Here, we used short-read metagenomic data and a database of publicly available isolate long-read sequencing to identify intragenic invertons in prokaryotes. In addition to using an existing short-read inverton calling program, we developed a long-read inverton finding pipeline, PhaVa, to more sensitively enumerate invertons. We identified intragenic invertons across the prokaryotic tree of life, in both bacteria and the small number of archaea that we evaluated. Using BTh as a model organism, we experimentally validated 10 intragenic invertons identified from our short-read metagenomic analysis. We further assessed the consequence of inversion by characterizing the phenotypic effects of an intragenic inverton found in the BTh thiamine biosynthesis gene *thiC*. Thiamine is an essential cofactor for many central metabolic processes and is bio-energetically costly to produce. Many microbes encode salvage, transport, and biosynthetic pathways ^51^. In BTh, thiamine acquisition and biosynthesis is highly regulated at both the transcriptional and translational level ^43,52^. Here, we find that thiamine biosynthesis also appears to be regulated at the genomic structural level. The *thiC* intragenic inverton induces a premature stop codon and we found that the truncated ‘reverse’ isoform has impaired growth in thiamine-limited conditions. However, we also found that at physiological concentrations of thiamine found in the human intestinal lumen, organisms encoding a locked ‘reverse’ isoform of *thiC* have a competitive growth advantage over the locked ‘forward’ isoform. This supports the presence of a novel mechanism of thiamine biosynthesis regulation and suggests a possible ecological explanation for the existence of a ‘toggle-able’ switch of isoforms.

While the advantages of each identified intragenic inverton will differ depending on the coding region affected, there are three general biological consequences of phase variation. One reason an organism may have preprogrammed heterogeneity is to enable division of labor. By generating subgroups within a population, members of the community may produce public goods at a potential cost to their own fitness but for the benefit of the group as a whole ^53–56^. While altruism in bacterial interspecies relationships is often unstable, as cheaters will take advantage of the public goods and outcompete, intraspecies altruism could bypass this as the losers in this scenario can be repopulated by the winners ^57^. The second type of heterogeneity producing behavior is via a bet hedging strategy ^58^. Diverse subgroups are generated allowing for survival in future selection events. One classic example of this is the CPS switching that occurs in many gut bacteria. As different CPS have varying susceptibility to phage predation, a diversified population allows the species to persist in the presence of phage ^27^. Third, bacteria of a given species and strain may exist in various biogeographic ‘niches’, where neighboring bacteria, host cells, nutrient access, and stressors might vary - thus, within a given ecological system such as the intestinal lumen, different subcommunities of bacteria may benefit from employing different bioregulatory programs. Future mechanistic work is needed to determine the advantages and community structure implications of each intragenic inverton.

Although we validated the presence of the intragenic invertons in BTth, we have not identified which invertases are responsible for each of the validated intragenic invertons that we described. We suspect that an underlying “molecular grammar” exists and that certain invertases recognize and flip specific sequences; specificity of invertases for a given sequence likely lie in the inverted repeats, but might also lie within the inverted regions. In terms of how the expression of these invertases is controlled and regulated in bacteria, phenotypic diversity is often generated via two different mechanisms; random ^57^ or coordinated specialization ^59^. It is possible that invertases function at a basal level and therefore there is a baseline, low level of inversion that occurs in a small proportion of the population. Alternatively, invertases may be expressed in response to specific cues or signals. As BTh encodes 56 invertases, future work is needed to identify which invertase flips these invertons and under which conditions. Understanding how these elements are regulated and the consequences of inversion could advance the field of synthetic biology and create new therapeutic targets.

Intragenic invertons that cause recoding mutations present an exciting opportunity to rethink gene variation. In BTh, we molecularly confirmed 2 recoding intragenic invertons (Table 1). Of note, these recoding mutations may help regulate the outer membrane of a bacterial cell potentially altering interactions with the host or other microbes. The first is BT0375, the putative CPS1 invertase. Cross regulation of CPS loci has been well described ^60,61^. Changes to the invertase structure could change its binding specificity and alter which regions it flips or its kinetics. Future studies are needed to elucidate how this inversion may add another layer of regulation to which CPS loci are expressed and when they are expressed. The second is *nagA* which encodes an enzyme that catalyzes the deacetylation of N-acetylglucosamine-6-phosphate (GlcNAc-6-P) to glucosamine-6-phosphate (GlcN-6-P). NagA is important for cell wall recycling and can supply GlcN-6-P for glycolysis ^62^. As NagA can be allosterically regulated ^63^, intragenic inversion could alter the binding of its allosteric regulator, altering this process; this might result in changing the metabolism of the cell or the outer membrane structure. Bacterial outer membranes play crucial roles in microbial interactions, niche establishment, and immune modulation. Intragenic invertons may add another layer of regulation and future studies are needed to study their effects.

While we find fairly extensive evidence of intragenic invertons using sequencing based approaches and explore some of them in detail, this work has limitations. First, our analysis of invertons across the prokaryotic tree of life was performed on previously sequenced isolates. While growth conditions for most of these samples are not readily available, we presume that most of these isolates were grown in rich laboratory media; these nutrient- and micronutrient-replete conditions may not recapitulate physiological conditions in which invertases are active or reverse orientations are favorable. However, there are currently limited long-read datasets available from physiological conditions for a wide range of prokaryotic organisms. We therefore may have only identified a minimal set of invertons in this study and we estimate the full “invertome” likely includes a larger number of elements. Second, if the invertons we identified are not representative of the true capacity for inversion, our gene set enrichment analysis may also not identify the types of genes hit most frequently in physiological conditions. Third, both PhaVa and PhaseFinder are reliant upon a reference genome or *de novo* assembly for read mapping and selection of a particular sequence for read mapping can affect inverton discovery. Detection of invertons is thus restricted to the genomic sequence common between the input sequenced strain and the reference. Finally, PhaVa uses relatively strict mapping parameters and if the selected reference is distantly related to the sequenced strain, read mapping quality will decrease and reduce the discovery rate. However, using a *de novo* assembly instead may result in missing ‘fully inverted’ invertons relative to reference strains, which may be of interest.

Despite these limitations, intragenic invertons are an exciting new mechanism for genetic variation and adaptation in bacteria. In this manuscript, we present a ‘roadmap’ for more in depth investigation of a specific invertible intragenic locus. Our initial analysis of long-read isolate data provides a minimal set of invertons, including intragenic, intergenic, and partial intergenic invertons. We expect future niche-specific investigation of inverton-containing organisms to identify additional invertons. Additionally, we anticipate that future studies of intragenic inverton will uncover new layers of bioregulation in prokaryotes, and more thoroughly demonstrate the many hidden genetic programs that exist within highly plastic bacterial genomes. More thorough characterization of invertons and other reversible and preprogrammed types of genomic variation will likely substantially impact several fields of research ranging from synthetic biology, to microbe-microbe interactions, to microbial physiology, and beyond.

## Materials and Methods

### Strains and media

The bacterial strains used in this study are listed in Table S1. *E. coli* strains were routinely grown in LB Miller media (Fisher) at 37 ℃. When necessary, carbenicillin was added at 100 µg/mL. BTh was grown anaerobically (90% Nitrogen, 5% carbon dioxide, 5% hydrogen) in an anaerobic chamber (Sheldon Manufacturing) in hemin (5 µg/mL) and L-cysteine (1 mg/mL) supplemented Brain Heart Infusion (Sigma) media (BHIS) or Varel-Bryant broth (VB)^64^. When necessary, the antibiotics tetracycline (2.5 µg/mL), erythromycin (25 µg/mL), or gentamicin (200 µg/mL) were added to the media. Thiamine HCL (Sigma) was added at the specified concentrations. All media used to grow BTh was preincubated in the anaerobic chamber overnight.

### Construction of *thiC* clean deletion and locked strains

The *thiC* clean deletion and locked strains were generated via allelic exchange as previously described ^65^. For Δ*thiC*, 600 - 700 base pair flanking regions of the coding region were amplified using Q5 high fidelity polymerase (New England Biolabs). Recombinant DNA used in this study is listed in Table S2. For locked strains, plasmid overhangs, flanking regions, locked repeats and intervening forward or inverted sequences were synthesized (Twist Biosciences) (Data S8). Regions were assembled into pExchange-tdk using the HiFi DNA Assembly Kit (New England Biolab). Plasmid inserts were verified using Sanger sequencing (ElimBio). Sequence confirmed plasmids were propagated in *E. coli* DH5α λ*pir*. *E. coli* S17-1 λ*pir* was used as a donor strain for conjugation into BTh Δ*tdk*. Exconjugants with chromosomal integration of plasmids were recovered on BHIS plates containing gentamicin and erythromycin. Second crossover events were selected by using BHIS FudR (200 µg/mL 5-fluoro-2-deoxy-uridine). Deletion and locked versions were confirmed by PCR.

To generate differentially resistant *thiC* locked strains, the suicide vectors pNBU2_tet and pNBU2_erm were used. *E. coli* S17-1 λ*pir* harboring the plasmids were used as donor strains for conjugation. Single crossover events were selected by plating on gentamicin plates containing either erythromycin or tetracycline respectively.

### Validating inversion in DNA

Intragenic inverton confirmation primers were designed using NCBI Primer Blast under default settings with the addition of adding in a GC clamp. PCR product size was targeted to be between 300 and 600 base pairs. The common and reverse primer were oriented on the same strand of the reference genome and the forward primer was located on the complementary strand. The common primer was located in between the two inverted repeats (fig. S2, primers listed in Data S6). Four of the predicted invertons were not experimentally tested, as target-specific primers could not be generated within the above constraints.

DNA was isolated from wild-type BTh cultures grown for 18 hours in either BHIS or VB media. DNA was isolated using a chemical and enzymatic lysis. Glass beads (0.1 mm) were added to bacterial pellets along with 700 µl of extraction buffer (50 mM Tris pH 7.5, 1 mM EDTA, 100 mM NaCl, 1% (w/v) SDS) and 25 µl of Proteinase K (10 mg/mL). Pellets were vortexed for 20 seconds and incubated at 55°C for 60 min. 700 µl of phenol:chloroform:isoamyl alcohol (25:24:1 by volume) was added to the mixture prior to incubating at room temperature for 5 minutes. Phases were separated by centrifuge at 10,000 rpm for 5 minutes. The aqueous upper layer was collected and transferred to a new tube. 5 µl of RNAse A (10 mg/mL) was added and incubated at 37°C for 15 minutes. An equal volume of phenol:chloroform:isoamyl alcohol was added, mixed, and incubated at room temperature for 5 minutes. Phases were separated as above and the aqueous phase was added to a new tube containing an equal volume of chloroform: isoamyl alcohol (24:1 by volume). Tubes were mixed and incubated at room temperature for 5 minutes prior to phase separation via centrifugation. The aqueous phase was added to a new tube along with 45 µl of 3M sodium acetate and 1 mL cold 100% ethanol. DNA was precipitated overnight at −20°C . Pellets were washed twice with 1mL of cold 70% ethanol. Dried pellets were resuspended in water.

PCR reactions were performed using Q5 high fidelity polymerase (68 ℃ annealing temp, 10 second annealing time, and 30 second extension time). PCR reactions were run on an 1.2% agarose gel. If multiple bands were visible, bands of the expected size were gel purified using the Qiagen Gel Extraction kit. If a single band of the expected size was observed, the PCR reaction was purified using the Monarch PCR Cleanup Kit (New England Biolabs). DNA was sent for Sanger sequencing. Sequencing was compared to the *in silico* prediction.

### Validating *thiC* inversion in RNA

RNA was isolated from wild-type BTh cultures grown for 18 hours in BHIS media. 5mL cultures were quenched using 500 µL phenol/ethanol solution (90% [vol/vol] ethanol and 10% [vol/vol] saturated phenol pH 4-5). Pellets were spun down and stored at −80 ℃ until extraction. Pellets were lysed in 250 µL PBS and 10 µL of lysozyme (10 mg/mL) at 37 ℃ for 30 minutes. 30 µl 20% SDS was added prior to an additional 30 minute incubation. 1.5 mL Trizol was added to the mixture and incubated at room temperature for 10 minutes. Chloroform (0.5 mL) was added to each sample and inverted vigorously for 15 seconds. The aqueous phase was taken from centrifuged samples and an equal volume of 100% ethanol was added. RNA was purified using the Zymo RNA clean kit. DNA was removed using Ambion Turbo DNAse. cDNA was made using Taqman Reverse Transcription reagents (Invitrogen) according to the manual. A no reverse transcriptase control was performed to ensure that all DNA was removed. PCR was performed to determine orientation of inverton as above. Correctly sized bands were sent for Sanger sequencing.

### BTh growth in thiamine concentrations

BTh wild-type, *thiC* locked forward, *thiC* locked reverse, and *thiC* knockout strains were grown overnight in BHIS media. Aliquots of each were then washed twice in preincubated PBS containing cysteine (1 mg/mL). Strains were inoculated at an OD600 of 0.05 in VB media containing the indicated concentration of Thiamine in a 96-well flat bottom plate. Readings were taken in a Stratus plate reader (Cerillo) every ten minutes. Non-inoculated VB media from each time point was used as a blank. The maximum OD600 value achieved per well was determined.

### Competitive growth assay

Marked BT0650 locked strains were grown overnight in BHIS with appropriate antibiotics. Strains were washed twice with preincubated PBS containing cysteine (1 mg/mL). A glass dilution tube containing 3 mL of VB with indicated concentrations of thiamine was inoculated with 1 x 10^3^ CFU/mL of each strain. After 40 hrs of growth at 37 ℃ in the anaerobic chamber, CFU/mL was determined by plating on selective agar. A competitive index was calculated by dividing the recovered CFU/mL of the locked forward by the CFU/mL of the locked reverse strain corrected by the inoculum.

### Identifying invertons in BTh with PhaseFinder

Two short-read datasets were used for identifying invertons in BTh, 416 samples from 149 adult HMT patients (^23^, BioProject PRJNA707487) and 142 samples from 21 pediatric HMT patients (^22^, BioProject PRJNA787952). Each individual short-read dataset was analyzed with PhaseFinder ^15^ with the VPI-5482 reference genome and default parameters to identify putative invertons in BTh. Invertons were included in further analysis if they had at least 5 reads mapping to the reverse orientation of the inverton, and had reads mapping to the reverse orientation in at least three different samples. Inverton-gene overlaps and partial overlaps were found using a custom script, now incorporated in PhaVa in the ‘Create’ step, and the gene annotations from the VPI-5482 genbank file (.gbff).

### The PhaVa algorithm

Inverted repeats are identified with einverted, part of the EMBOSS suite ^66^. For each putative inverton, two sequences are then created: one where the sequence between identified inverted repeat pairs is inverted (reverse) and one where it is not (forward), along with flanking sequence on either side, similar to PhaseFinder. Long-reads are mapped against the created sequence with minimap2 ^67^ and must pass several filters to be included as evidence of inversion. 1) reads must have a MAPQ score of ≥ 2 to eliminate multimapping reads. 2) Reads must span the entire length of the inverton and at least 30 bps into the flanking sequence on either side. 3) The mismatch rate along the length of a read must be below a maximum mismatch rate. The mismatch rate is considered separately over the length of an inverton and over flanking sequence, to avoid reads that map well to only one region or the other. An adjustable mismatch rate is used instead of a flat mismatch cutoff to account for both the variable length of long-reads and the high sequencing error rate of current long-read sequencing technologies relative to short-read sequencing. After mapping, reads mapped to the inverted and non-inverted sequences are tallied and optional post-mapping filters are applied. 1) A minimum number of total reads mapped to the ‘reverse’ sequence and 2) a minimum proportion of total mapped reads mapped to the ‘reverse’ sequence.

### Simulating long-read datasets for optimizing PhaVa

For benchmarking, ten bacterial species were selected, in part based on the relevance in the human microbiome. For each species, a reference genome and reference long-read dataset were obtained from NCBI (File S3). Long-reads were mapped against their respective reference genome with minimap2 and the mappings were used as input for the ‘characterization stage’ of NanoSim ^68^ in genome mode. The resulting NanoSim models were used to generate simulated long-read datasets in the ‘simulation stage’ in genome mode. Reads were generated from the unmodified reference genome, and so no evidence of inversion for any inverton is expected, and any invertons identified by PhaVa would be false positives. For each species, five coverage levels and six replicates of each coverage level were generated (Data S7) totaling in 300 simulated long-read datasets. *Streptomyces eurocidicus* and *Enterococcus faecalis* simulated readsets were generated at relatively deeper coverages due to poor read mapping characteristics from the selected reference long-read datasets resulting in a smaller proportion of reads passing PhaVa read mapping filters. Simulated readsets were then analyzed with PhaVa and used to estimate false positive rates and optimize post-mapping filters.

### Identifying putative invertons from public long-read sequencing data with PhaVa

Candidate isolate long-read sequencing datasets were identified on NCBI with the following search criteria: “(Bacteria[Organism] OR Archaea[Organism]) AND ("pacbio smrt"[Platform] OR "oxford nanopore"[Platform]) AND genomic[Source]”. Datasets were further filtered by removing datasets with the “amplicon” flag, and removing datasets with less than 50 Mbp of sequencing in total (Data S5). Individual read datasets were downloaded with fastq-dump, a part of the sratoolkit (https://trace.ncbi.nlm.nih.gov/Traces/sra/sra.cgi?view=software). Nanostat ^69^ was run on each remaining readset to measure dataset characteristics. For each unique taxid represented in the resulting readsets, a reference genome and paired genbank file (.gbff) were selected by identifying a genome with the highest level of completion for that species, and the least number of contigs. In the case of reference genomes with equal quality based on these parameters, the first identified was selected. Long-read datasets were then analyzed with PhaVa with default parameters. Gene overlaps and partial gene overlaps were identified by comparing coordinates of genbank file annotations with inverton coordinates, a function available for use in PhaVa (fig. S8).

### AlphaFold prediction

Structural predictions of the amino acid sequences for the forward and reverse orientations of the intragenic inverton within BT0375 were generated using AlphaFold ^70^ v2.2.0. The required databases were downloaded on March 3rd, 2022 and the max template date was set to 2020-05-14. The top ranked structures were then visualized and aligned in PyMOL.

### Gene set enrichment analysis

In order to assess which functional groups were enriched for genes harboring intragenic invertons, we performed a clade-resolved gene set enrichment analysis. We first annotated genes with KEGG KOs using the kofamKOALA tool ^71^ and with Pfam domains by running HMMER3 ^72^ with the Pfam domain database. KEGG pathways and modules were filtered for those that were present in bacterial genomes and Pfam clan definitions were downloaded from the Interpro website ^73^. We then calculated enrichments per genome and additionally per species and per genus, for those combining the genes from all genomes in a specific clade. At each level, we filtered out groups with fewer than 5 intragenic invertons (fig. S5), resulting in 10 genomes, 12 species, and 19 genera being included for downstream analysis. Alternatively, we also considered genes with both intragenic or partial intergenic invertons, resulting in 47 genomes, 52 species, and 54 genera being tested. In each group, we tested for each pathway if the genes annotated with this pathway were enriched for those carrying invertons by using a one-sided Fisher test. Pathways, the genes of which did not harbor any invertons in a specific group, were skipped for the enrichment analysis for a given group. Multiple testing correction was performed with the Benjamini-Hochberg procedure ^74^.

### Identifying putative invertons from long-read metagenomes

200 hybrid short-read and long-read human stool metagenomic datasets were accessed from BioProject PRJNA820119 ^38^. Each hybrid dataset was assembled using SPAdes ^75^ with the ‘-meta’ flag and long-reads provided with the ‘--nanopore’ option. An additional ten nanopore long-read human stool microbiome metagenomic datasets from BioProject PRJNA940499 ^39^ were assembled with Flye ^76^. Assembled contigs will be deposited at https://doi.org/10.5281/zenodo.7662825 after publication. Gene annotations for assemblies were obtained with Prodigal ^33^ using the ‘-meta’ flag. Contig taxonomic classifications were obtained with Kraken2 ^77^. Each long-read dataset was then analyzed with PhaVa with default parameters, using its respective de novo assembly as its reference assembly. Resulting inverton calls were dereplicated by clustering the inverton with 1000 bp flanking sequence upstream and downstream at 99% average nucleotide identity with CD-HIT ^78^.

## Supplemental Figures

**Fig. S1.**
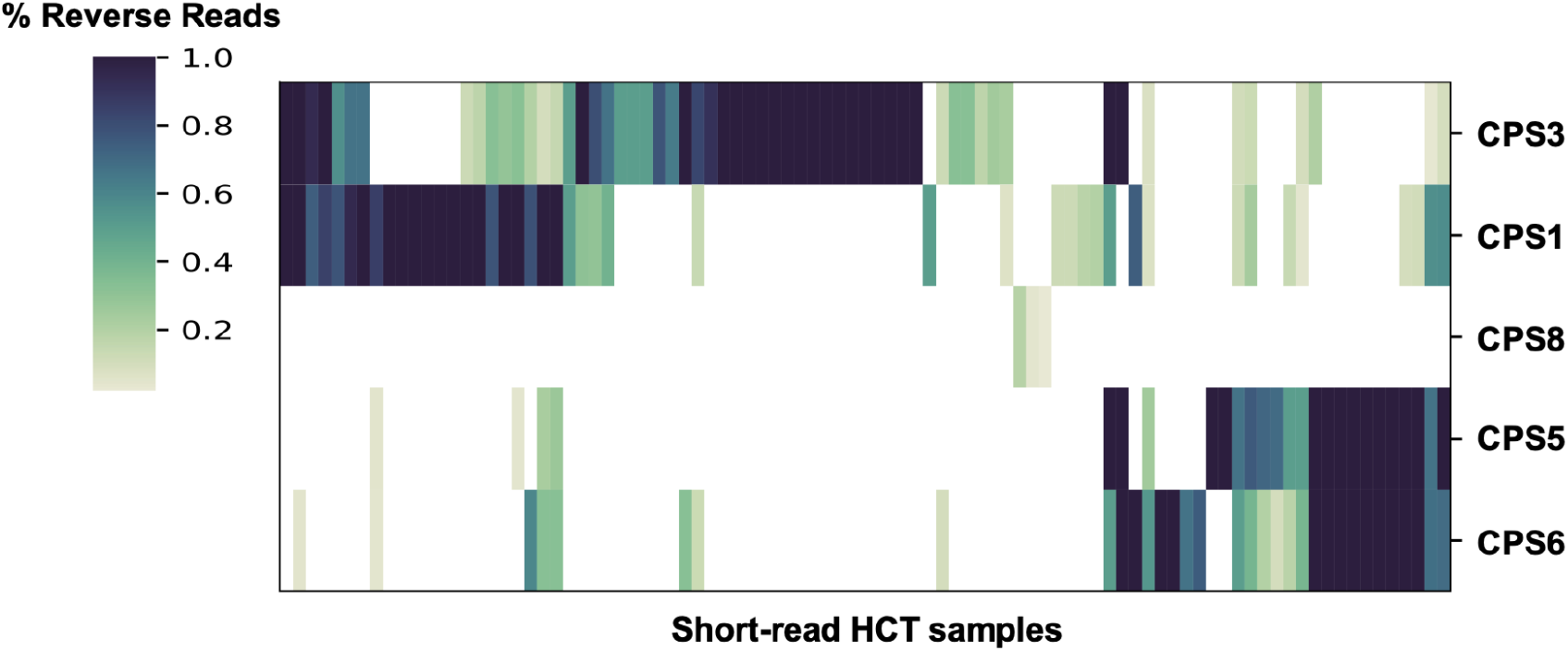
Inversion proportion of CPS loci invertons in BTh. Inversion proportions of CPS loci invertons in HCT metagenomic samples measured with PhaseFinder. Samples with no inversions in the five CPS invertons were removed.

**Fig. S2.**
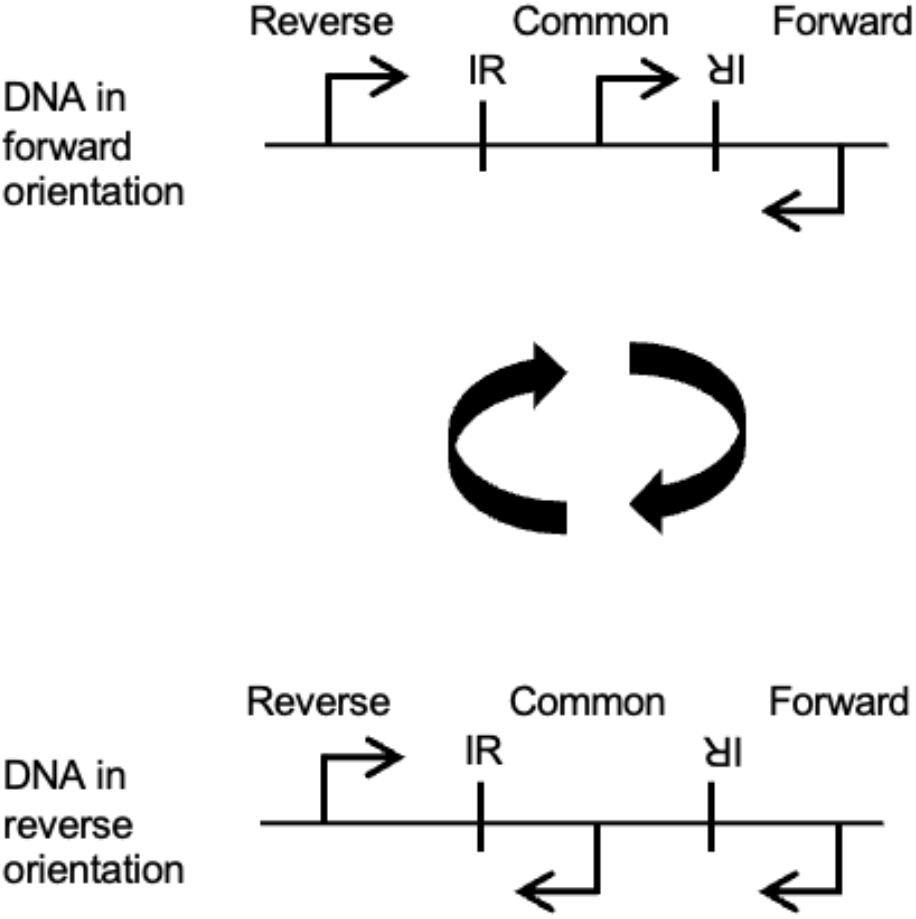
Inverton confirmation PCR primer design. A Forward and Reverse primer bind to regions of the genome upstream and downstream of the inverton on opposite strands. The Common primer binds the DNA inside of the inverton, between the inverted repeats. When the DNA is in the forward orientation, the Common and Forward primer will generate a PCR product. When the inverton flips, the Common and Reverse primer will generate a PCR product.

**Fig. S3.**
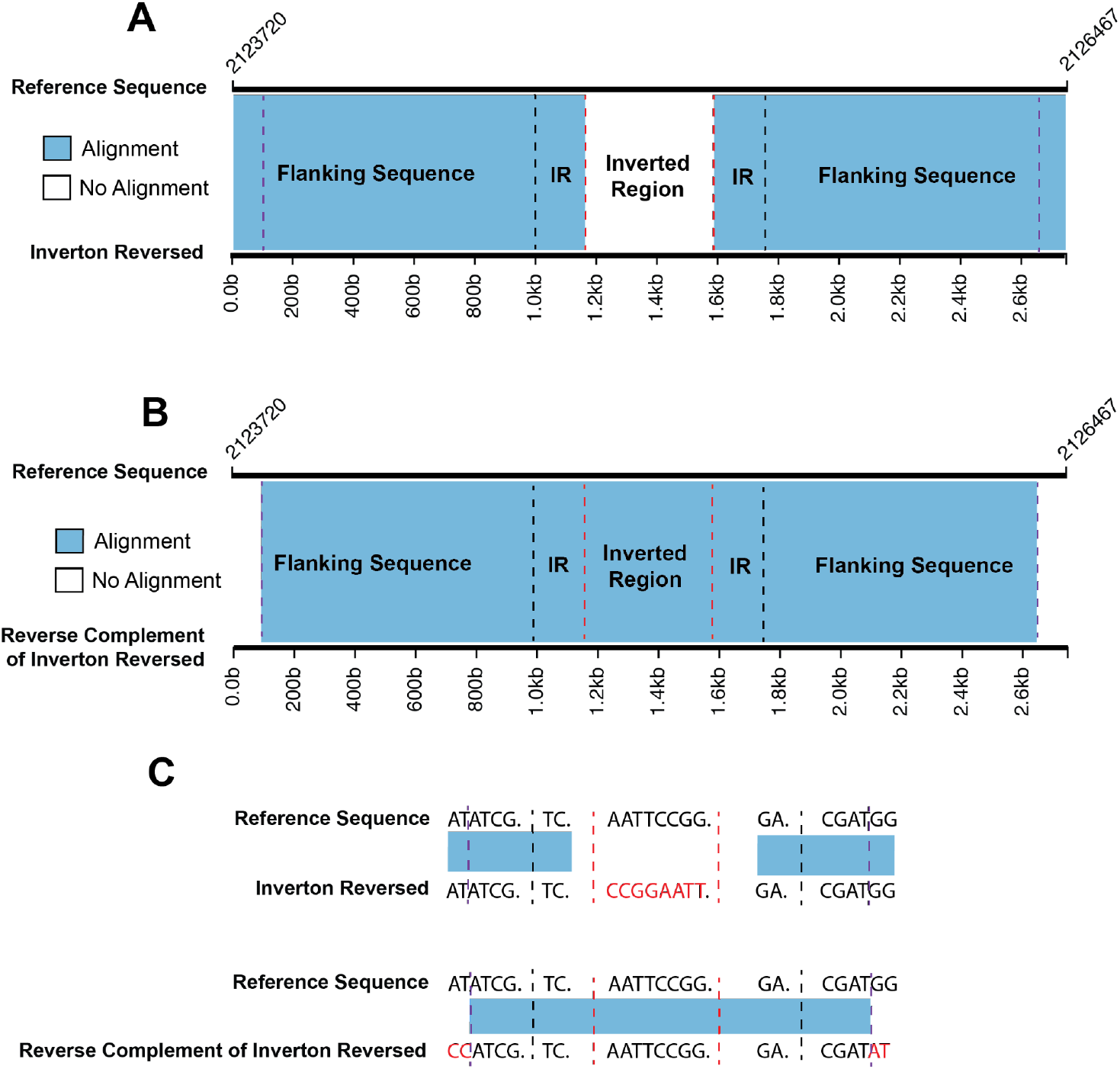
Very long (>750bp), near perfect, inverted repeats can lead to false positives. (**A**) Alignment of inverton NZ_CP025371.1:2124719-2124870-2125316-2125467, with its invertible sequence inverted, against the *B. pertussis* genome leads to perfect alignment of flanking and IR regions as expected. ‘Reference genome’ refers to the *B. pertussis* reference genome sequence. ‘Inverton reversed’ refers to the putative inverton sequence and flanking sequence, with the invertible sequence inverted. Red dashed lines indicate boundaries of the invertible sequence, black dashed lines indicate boundaries of the inverted repeats as detected by einverted, and purple dashed lines indicate the true boundary of inverted repeats. (**B**) Alignment of the reverse complement of the entire inverton NZ_CP025371.1:2124719-2124870-2125316-2125467 with its invertible sequence inverted and flanking sequence, against the *B. pertussis* genome leads to near perfect alignment (6 mismatches) spanning far into the flanking sequence to the true boundary of the inverted repeats, allowing for reads to map regardless of inverton orientation. (**C**) Example with toy nucleotide sequences. Red nucleotides indicate mismatches.

**Fig. S4.**
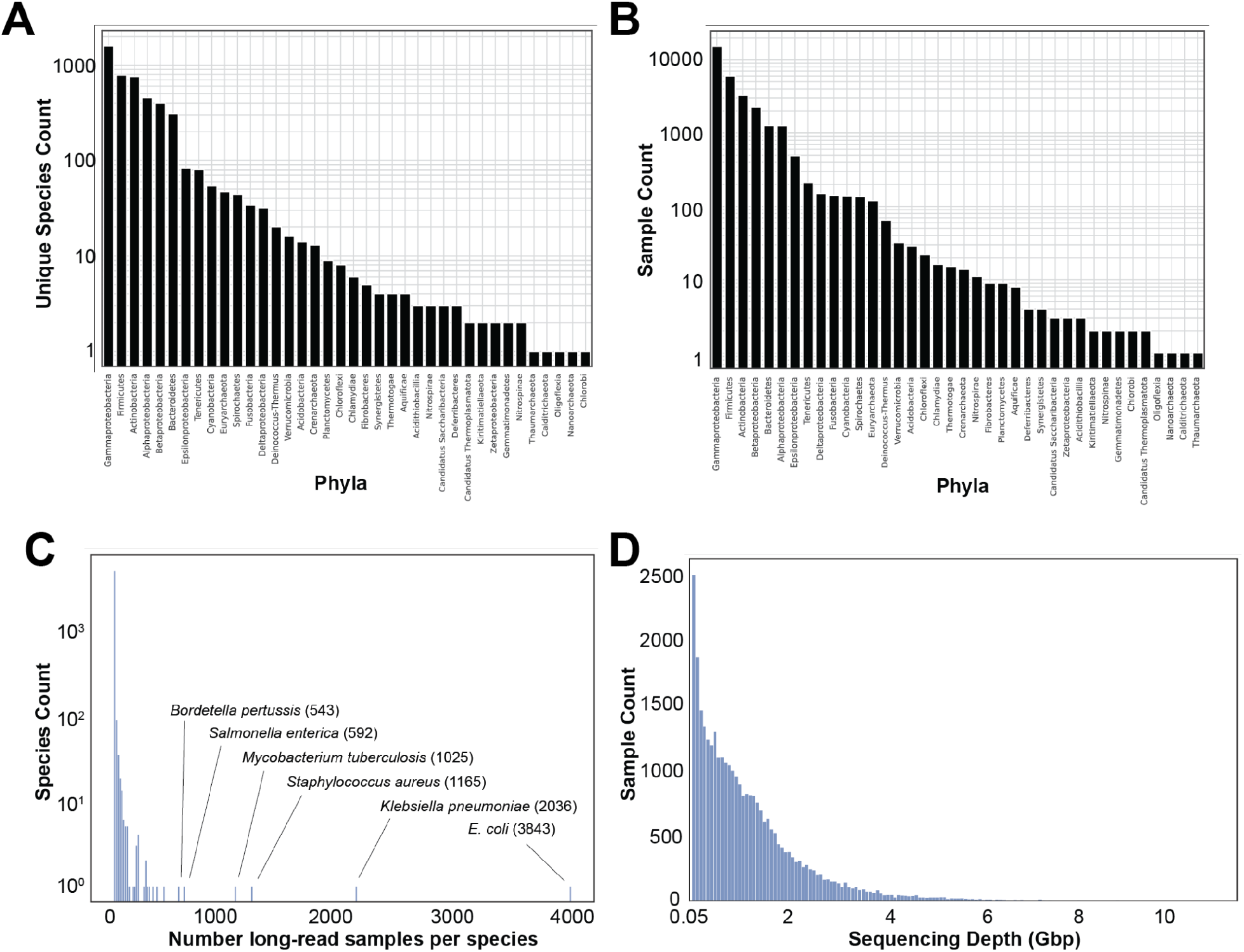
Overview of SRA long-read isolate sequencing samples analyzed with PhaVa. (**A**) The number of unique species represented in the dataset, grouped by phylum. (**B**) The raw number of sequencing samples, grouped by phylum. (**C**) Histogram of sequencing samples per species. Species with particularly large numbers of samples are labeled. (**D**) A histogram of sequencing depths for all long-read isolate sequencing samples.

**Fig. S5.**
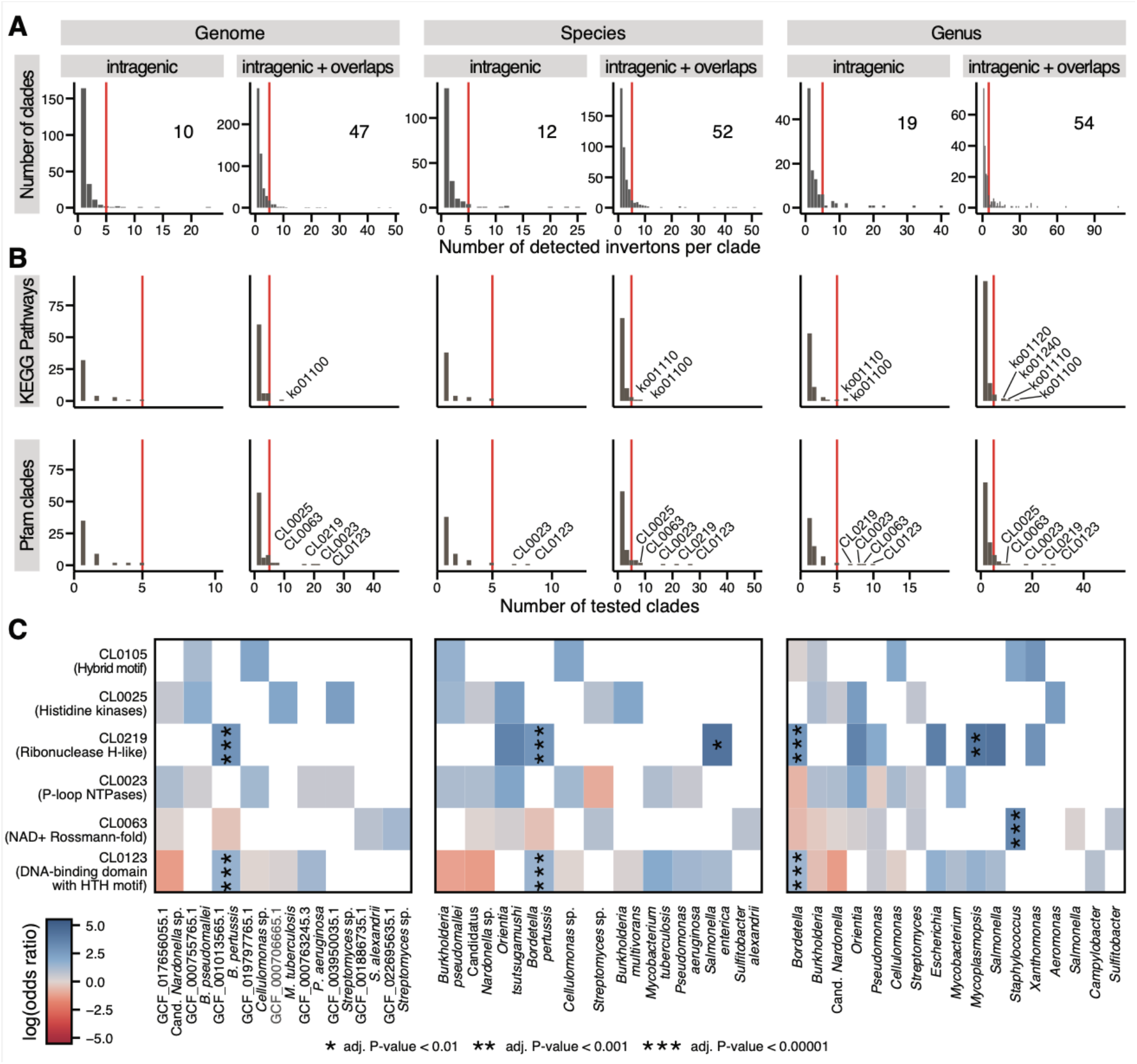
Intragenic invertons are rare across genomes yet consistently enriched in some Pfam clans. **(A)** Histograms showing the number of clades (genomes, species, or genera) at various numbers of invertons indicate that invertons are rare, as only one to three invertons can be detected in the majority of clades. Only clades with at least five invertons (red line; number of clades is indicated in the top-right corner of each subplot) were included for the subsequent enrichment analysis. **(B)** KEGG pathways and Pfam clans were tested for enrichment of intragenic (or partial intergenic) invertons in included clades, using a one-sided Fisher’s exact test per clade (see Methods). Enrichment was only calculated for sets with at least five invertons associated with genes in the set. Histograms show the number of sets with enrichment score at the number of included clades, showing that most enrichments could be calculated for single clades only. For example, all KEGG pathways associated with enough intragenic invertons for an enrichment analysis on genome-level were specific for each genome. Sets with enrichment scores across at least five clades (red line) are labeled with their corresponding identifiers. **(C)** Heatmap showing the log-odds ratio (effect size for the enrichment of intragenic invertons) across included clades for the six Pfam clans that have enrichment scores on genus-level (see panel B). Stars indicate significance of the enrichment as calculated by Fisher’s exact test and corrected for multiple hypothesis testing using the Benjamini-Hochberg procedure.

**Fig. S6.**
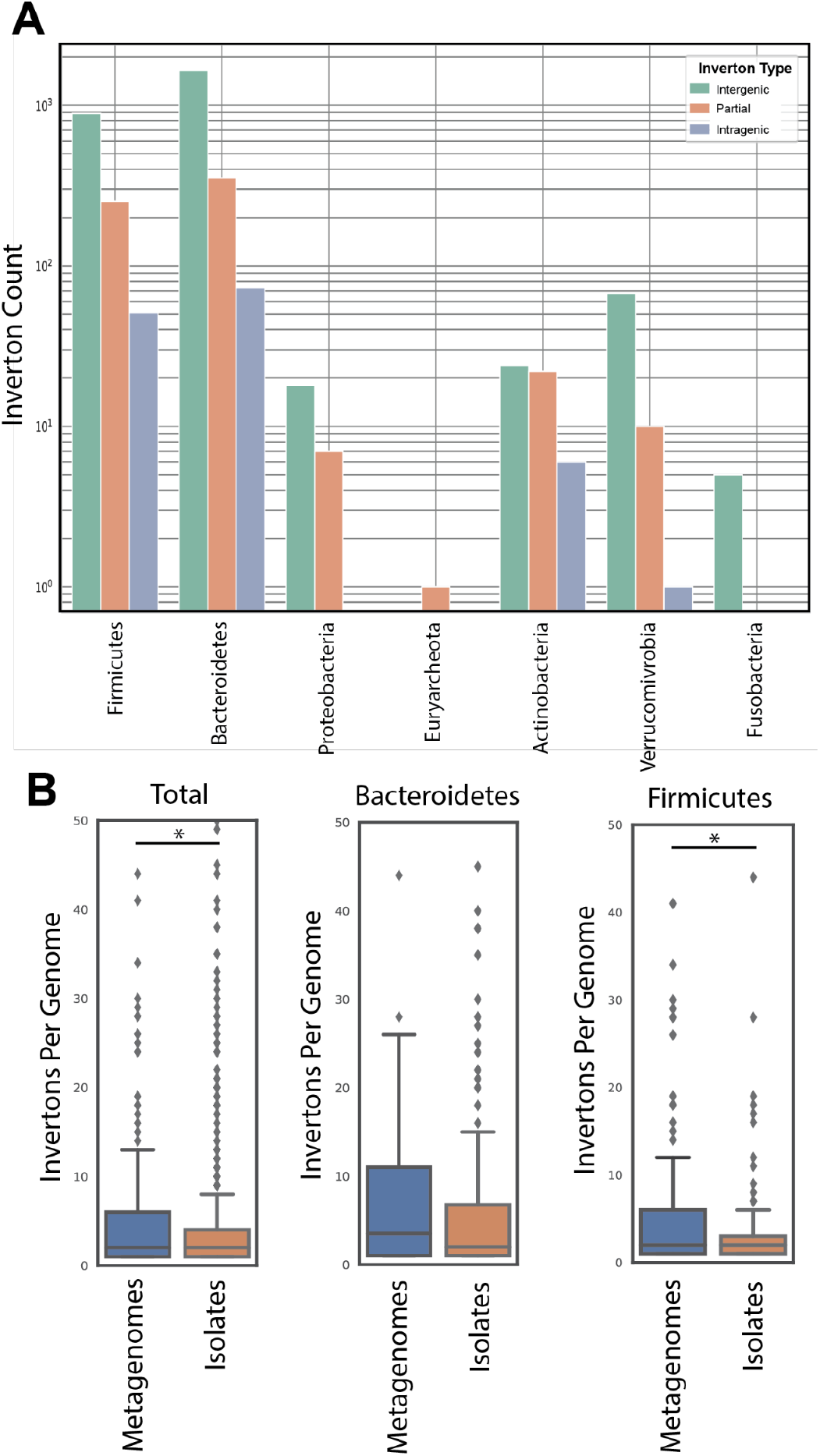
PhaVa analysis of 210 long-read metagenomes from human stool. (**A**) Counts of invertons identified with PhaVa in 210 stool samples, grouped by phylum and the type of inverton. (**B**) Comparisons of the number of invertons (per genome) found in metagenomic datasets vs. SRA isolate sequencing samples. Total refers to all invertons identified, regardless of taxonomic classification. The distribution of inverton counts per species were found to be significantly different between metagenomes and isolate samples in both the Total and Firmicutes comparisons (p=3.35e-05 and p=0.005 respectively) with a Kolmogorov–Smirnov test. Other individual phyla were not compared due to small species counts with invertons in metagenomic samples.

**Fig. S7.**
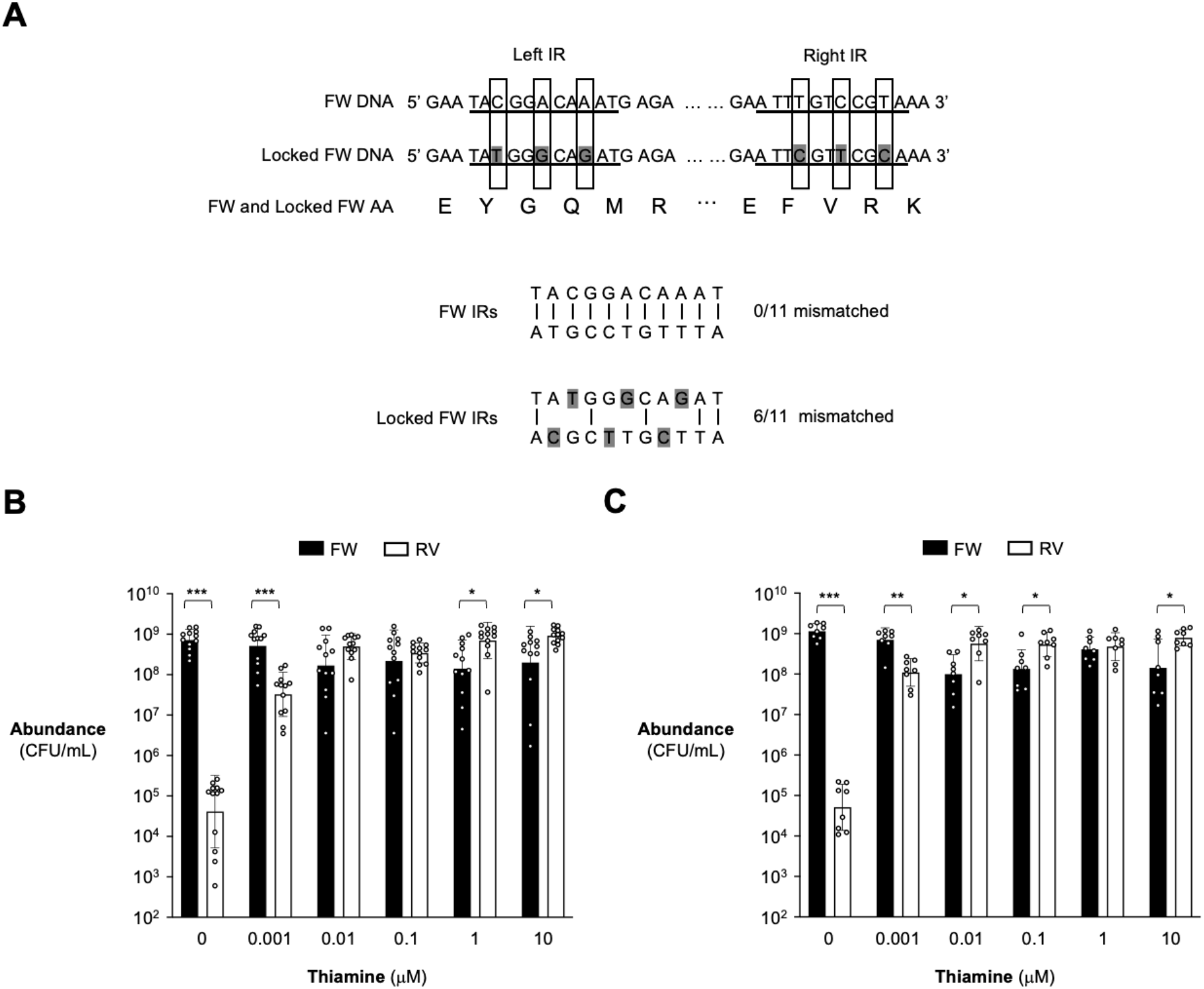
Locked *thiC* intragenic inverton construction and growth competition. (**A**) Generation of locked intragenic invertons. The forward and locked forward *thiC* IR nucleotide sequences are shown. When possible, the wobble position of each codon corresponding to the IR was mutated to increase mismatches between the two palindromic sequences while maintaining the amino acid sequence. Nucleotides that were mutated are highlighted in gray. (**B**-**C**) Locked *thiC* strains were competed against each other in thiamine-containing media in a 1:1 ratio. After 40 hours, the abundance of each strain was enumerated using selective agar. Black bars indicate the locked forward strain and white bars indicate the locked reverse strain. Recovered abundances shown here correspond with the competitive index shown in Fig. 4D. In (**B**) the locked forward strain is marked with an erythromycin resistant cassette and the locked reverse strain is marked with a tetracycline resistant cassette. In (**C**) the locked forward strain is marked with a tetracycline resistant cassette and the locked reverse strain is marked with an erythromycin resistant cassette. Geometric mean and geometric standard deviation are shown for replicates conducted across 4-6 independent experiments. For each thiamine concentration a ratio paired t test was performed on the locked forward and locked reverse abundances. ***, p < 0.001; **, p < 0.01; *, p < 0.05.

**Figure S8:**
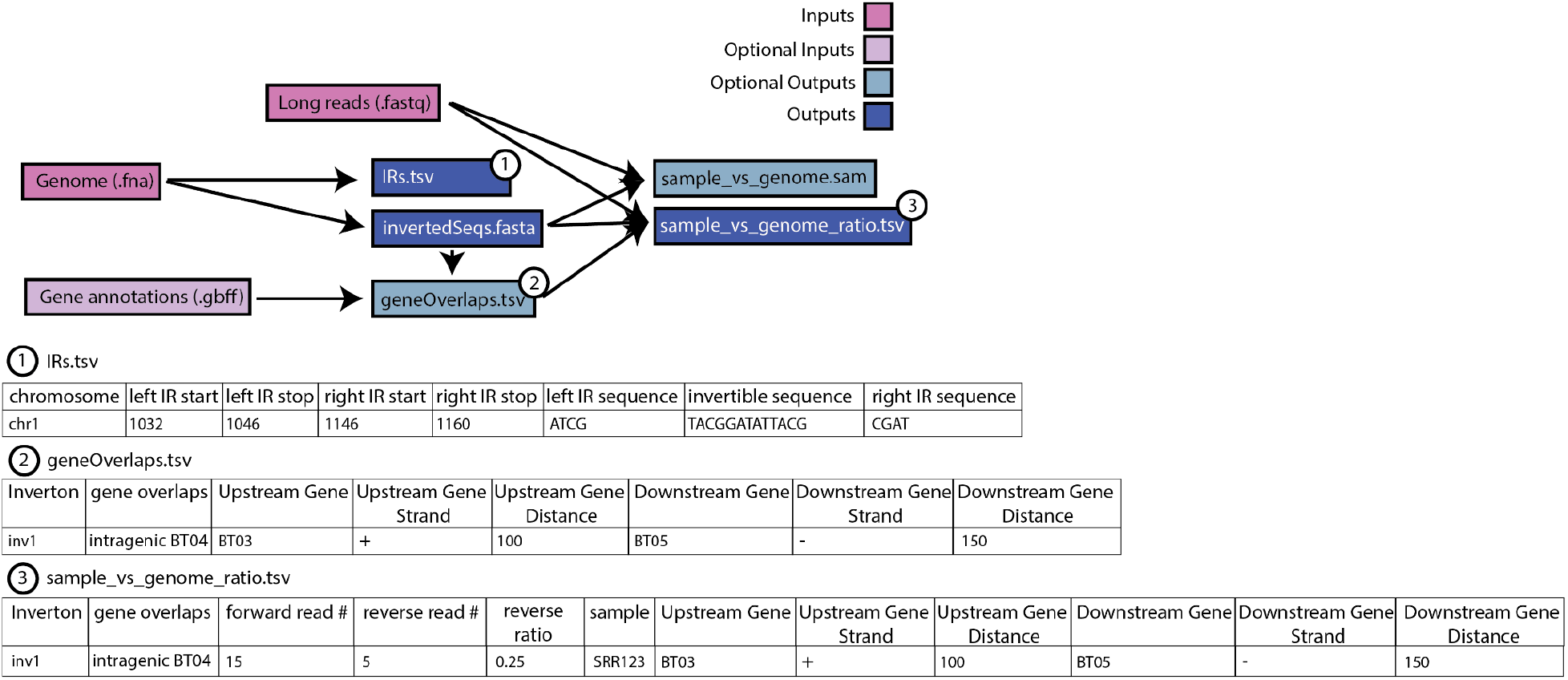
Inputs and outputs of a variation_wf PhaVa run. Output tables of particular interest are labeled and shown below the diagram with example output.

**Table S1.** Strains used in this study.

| Strain name | Source | Identifier |
| --- | --- | --- |
| *Bacteroides thetaiotaomicron* VPI-5482 $\Delta tdk$ | <sup>79</sup> | WT |
| *Bacteroides thetaiotaomicron* VPI-5482 $\Delta tdk$ $\Delta BT0650$ | this study | RC131 |
| *Bacteroides thetaiotaomicron* VPI-5482 $\Delta tdk$ BT0650 locked RV | this study | RC149 |
| *Bacteroides thetaiotaomicron* VPI-5482 $\Delta tdk$ BT0650 locked FW | this study | RC134 |
| *Bacteroides thetaiotaomicron* VPI-5482 $\Delta tdk$ BT0650 locked FW<br>*NBU2::NBU2_tet* | this study | RC165 |
| *Bacteroides thetaiotaomicron* VPI-5482 $\Delta tdk$ BT0650 locked FW<br>*NBU2::NBU2_erm* | this study | RC 166 |
| *Bacteroides thetaiotaomicron* VPI-5482 $\Delta tdk$ BT0650 locked RV<br>*NBU2::NBU2_erm* | this study | RC164 |
| *Bacteroides thetaiotaomicron* VPI-5482 $\Delta tdk$ BT0650 locked RV<br>*NBU2::NBU2_tet* | this study | RC163 |
| *E. coli* S17-1 $\lambda pir$; *zxx::RP4 2-(Tetr::Mu) (Kanr::Tn7) \lambda pir* | <sup>80</sup> | S17-1 $\lambda pir$ |
| *E. coli* DH5α $\lambda pir$; *F- endA1 hsdR17 (r-m+) supE44 thi-1 recA1 gyrA relA1 \Delta(lacZYA-argF)U189 \phi80lacZ\Delta M15 \lambda pir* | <sup>81</sup> | DH5α $\lambda pir$ |

**Table S2.** Recombinant DNA used in this study.

| Recombinant DNA | Identifier | Source |
| --- | --- | --- |
| pKNOCK-*bla-ermGb::tdk* | pExchange | 79 |
| pExchange BT0650 KO | pRBC20 | this study |
| pExchange BT0650 locked FW | pRBC21 | this study |
| pExchange BT0650 locked RV | pRBC22 | this study |
| pNBU2_tet | tetR | 24 |
| pNBU2_erm | ermR | 24 |

## Acknowledgments

We thank Nora Enright, Danica Schmidtke, Aravind Natarajan, Jack Diaz Shanahan, Dylan Maghini, Mai Dvorak, Alvin Han, Meena Chakraborty, and Bhatt Lab members for helpful conversations and scientific advice regarding this project. We also thank Wenhan Zhu for plasmids (pNBU2_tet and pNBU2_erm) and Sebastian Winter for the strain (DH5α) that were used in the context of this project. We also thank Daniel Haft and Francoise Thibaud-Nissen at NCBI for helpful discussion about accessing SRA long-read datasets.

## Funding

National Institutes of Health R01 AI148623 (ASB)

National Institutes of Health R01 AI143757 (ASB)

Stand Up 2 Cancer Foundation

National Institutes of Health T32 training Grant HG000044 (RBC)

National Institutes of Health T32 training Grant HL120824 (PTW)

## Author Contributions

Conceptualization: RBC, PTW, ASB

Methodology: RBC, PTW, JW

Investigation: RBC, PTW, JW, RMP, GZMG, ASH, MOG, EFB, AMM

Visualization: RBC, PTW, JW, RMP

Funding acquisition: ASB

Project administration: RBC, PTW, ASB

Supervision: RBC, PTW, ASB

Writing – original draft: RBC, PTW, ASB

Writing – review & editing: RBC, PTW, JW, RMP, GZMG, ASH, MOG, EFB, AMM, ASB

## Data and materials availability

PhaVa is available at (https://github.com/patrickwest/PhaVa). Short-read adult HCT stool sequencing data was previously published and is available at (NCBI BioProject ID PRJNA707487). Short-read pediatric HCT stool sequencing data was previously published and is available at (NCBI BioProject ID PRJNA787952). Long-read metagenomic sequencing data was previously published and is available at BioProject PRJNA820119 and BioProject PRJNA940499. Assembled metagenomic contigs will be made available after publication at https://doi.org/10.5281/zenodo.7662825. A list of accession numbers for long-read isolate sequencing data is available in supplementary file Data S5.

## Supplementary Information is available for this paper

### Extended Data Tables

Data S1. *B. theta* intragenic invertons identified from short-read metagenomic sequencing samples.

Data S2. Archaeal invertons identified from long-read isolate sequencing samples.

Data S3. Invertons identified from long-read isolate sequencing samples.

Data S4. Dereplicated invertons identified from long-read metagenomic sequencing samples.

Data S5. List of accession numbers and associated metadata for long-read isolate sequencing samples.

Data S6. Primers used in this study.

Data S7. Simulated read datasets.

Data S8. Additional sequences.

