## Supplementary material for "Intragenic DNA inversions expand bacterial coding capacity": Data S8

Sequences for locked BT0650 invertons

Lowercase letters - wings of homology for HiFi assembly

Underline - locked inverted repeats

BT0650 Locked FW insert

cggccgctctagaactagtgCCGGAATCGGTGCACCGAGCCACGCTGCCGAAGCAATGGAACTGGGTGCTTCTGCCGTACTGGTAAATACCGCTATCGCCGTAGCAGGCAATCCGGTAGAAATGGCGTTGGCCTTCAAAGCCGCCACCGAAGCCGGAAGACGGGCATACGAAGCAGGACTGGGTCTGCAGGCTGATAACTTTATAGCAGAAGCAAGTTCACCTTTAACTGCATTTTTGGAATAAGTAATAGGTGATAAGTAATAGGTAATAAGTAGGGAGTAGTGTTTGTTTCACTATTCTTAATGATATGATAATAAAATTCTATGGAACAAAAAATAAAATTCCCCCGTTCTCAGAAAGTGTATCTGCCGGGTAAACTCTATCCGAACATCCGTGTAGCCATGCGGAAAGTTGAACAAGTGCCCAGTGTCAGCTTCGAAGGAGAAGAAAAGATAGCTACTCCGAATCCGGAAGTATATGTATATGACACCAGCGGTCCGTTCAGTGACCCGTCAATGAGCATTGATCTTAAGAAAGGACTTCCCCGCTTACGTGAAGAATGGATTGTGGGACGCGGTGACGTAGAGCAACTCCCCGAAATCACTTCGGAATATGGGCAGATGAGACGTGATGACAAAAGTCTGGATCATCTCCGCTTTGAACACATTGCCCTTCCCTATCGTGCCAAAAAGGGAGAAGCCATCACTCAGATGGCATACGCCAAACGAGGAATCATCACTCCGGAAATGGAATATGTGGCGATCCGCGAAAATATGAACTGTGAGGAACTGGGAATCGAGACTCACATTACTCCCGAATTCGTTCGCAAAGAAATAGCAGAAGGACATGCAGTGCTGCCTGCCAACATCAATCATCCGGAAGCCGAACCGATGATTATCGGCCGTAACTTCCTAGTGAAAATCAATACGAATATCGGAAACTCTGCCACCACTTCCAGCATTGACGAGGAAGTGGAAAAAGCTTTATGGAGCTGCAAATGGGGAGGTGATACGCTGATGGACCTTTCAACAGGAGAAAATATTCATGAGACACGTGAATGGATCATCCGCAACTGCCCTGTACCGGTAGGCACTGTGCCTATCTATCAGGCACTTGAGAAAGTGAATGGCGTAGTGGAAGACCTTAACTGGGAAATCTATCGGGACACACTGATCGAACAATGCGAACAGGGAGTAGATTATTTTACGATTCATGCAGGTATCCGCCGTCATAATGTTCATCTTGCCGATAAACGTCTGTGCGGCATTGTCAGCCGTGGAGGAAGCATCATGAGTAAATGGTGTCTGGTGCACGATCAGGAAAGTTTCCTGTATGAACACTTTGACGATATCTGTGATATTCTGGCTCAATACGACGTTGCCGTATCATTGGGAGACGGACTGCGTCCGGGCTCTATCCccccgggctgcaggaattcg

BT0650 Locked RV insert

cggccgctctagaactagtgCCGGAATCGGTGCACCGAGCCACGCTGCCGAAGCAATGGAACTGGGTGCTTCTGCCGTACTGGTAAATACCGCTATCGCCGTAGCAGGCAATCCGGTAGAAATGGCGTTGGCCTTCAAAGCCGCCACCGAAGCCGGAAGACGGGCATACGAAGCAGGACTGGGTCTGCAGGCTGATAACTTTATAGCAGAAGCAAGTTCACCTTTAACTGCATTTTTGGAATAAGTAATAGGTGATAAGTAATAGGTAATAAGTAGGGAGTAGTGTTTGTTTCACTATTCTTAATGATATGATAATAAAATTCTATGGAACAAAAAATAAAATTCCCCCGTTCTCAGAAAGTGTATCTGCCGGGTAAACTCTATCCGAACATCCGTGTAGCCATGCGGAAAGTTGAACAAGTGCCCAGTGTCAGCTTCGAAGGAGAAGAAAAGATAGCTACTCCGAATCCGGAAGTATATGTATATGACACCAGCGGTCCGTTCAGTGACCCGTCAATGAGCATTGATCTTAAGAAAGGACTTCCCCGCTTACGTGAAGAATGGATTGTGGGACGCGGTGACGTAGAGCAACTCCCCGAAATCACTTCGGAATATGGGCAGATTCGGGAGTAATGTGAGTCTCGATTCCCAGTTCCTCACAGTTCATATTTTCGCGGATCGCCACATATTCCATTTCCGGAGTGATGATTCCTCGTTTGGCGTATGCCATCTGAGTGATGGCTTCTCCCTTTTTGGCACGATAGGGAAGGGCAATGTGTTCAAAGCGGAGATGATCCAGACTTTTGTCATCACGTCTCATTCGTTCGCAAAGAAATAGCAGAAGGACATGCAGTGCTGCCTGCCAACATCAATCATCCGGAAGCCGAACCGATGATTATCGGCCGTAACTTCCTAGTGAAAATCAATACGAATATCGGAAACTCTGCCACCACTTCCAGCATTGACGAGGAAGTGGAAAAAGCTTTATGGAGCTGCAAATGGGGAGGTGATACGCTGATGGACCTTTCAACAGGAGAAAATATTCATGAGACACGTGAATGGATCATCCGCAACTGCCCTGTACCGGTAGGCACTGTGCCTATCTATCAGGCACTTGAGAAAGTGAATGGCGTAGTGGAAGACCTTAACTGGGAAATCTATCGGGACACACTGATCGAACAATGCGAACAGGGAGTAGATTATTTTACGATTCATGCAGGTATCCGCCGTCATAATGTTCATCTTGCCGATAAACGTCTGTGCGGCATTGTCAGCCGTGGAGGAAGCATCATGAGTAAATGGTGTCTGGTGCACGATCAGGAAAGTTTCCTGTATGAACACTTTGACGATATCTGTGATATTCTGGCTCAATACGACGTTGCCGTATCATTGGGAGACGGACTGCGTCCGGGCTCTATCCccccgggctgcaggaattcg

Amino acid sequences for BT0375

Forward

VESGRAPYVHRLFHDVYTGVDICQKKALPVGELNRLLYEDPKSERLRRTQAIAALMFQFCGMSFADLAHLEKSSLERNIIRYNRIKTKTPMSVEVLDTAQDIISRLRNCQPSHPDCPDYLFSILLGDKKREDESAYREYQSALRRFNNRLKRLAKALRLTSPVTSYTIRHSWATTAKYRGVPIEMISESLGHKSIKTTQIYLKGFGLQERTEVNRMNLSYVKNCRIGRV

Reverse

VESGRAPYVHRLFHDVYTGVDICQKKALPVGELNRLLYEDPKSERLRRTQAIAALMFQFCGMSFADLAHLEKSSLERNIIRYNRIKTKTPMSVEVLDTAQIVGTIRMRRLTVAKPADYILFSILLGDKKREDESAYREYQSALRRFNNRLKRLAKALRLTSPVTSYTIRHSWATTAKYRGVPIEMISESLGHKSIKTTQIYLKGFGLQERTEVNRMNLSYVKNCRIGRV
